# ENHANCEMENT OF PARVALBUMIN INTERNEURON-MEDIATED NEUROTRANSMISSION IN THE RETROSPLENIAL CORTEX OF ADOLESCENT MICE FOLLOWING THIRD TRIMESTER-EQUIVALENT ETHANOL EXPOSURE

**DOI:** 10.1101/2020.06.16.155077

**Authors:** Clark W. Bird, Glenna J. Chavez, Megan J. Barber, C. Fernando Valenzuela

**Affiliations:** Department of Neurosciences, School of Medicine, University of New Mexico Health Sciences Center Albuquerque, New Mexico, USA

## Abstract

Prenatal ethanol exposure causes a variety of cognitive deficits that have a persistent impact on quality of life, some of which may be explained by ethanol-induced alterations in interneuron function. Studies from several laboratories, including our own, have demonstrated that a single binge-like ethanol exposure during the equivalent to the third trimester of human pregnancy leads to acute apoptosis and long-term loss of interneurons in the rodent retrosplenial cortex (RSC). The RSC is interconnected with the hippocampus, thalamus, and other neocortical regions and plays distinct roles in visuospatial processing and storage, as well as retrieval of hippocampal-dependent episodic memories. Here we used slice electrophysiology to characterize the acute effects of ethanol on GABAergic neurotransmission in the RSC of neonatal mice, as well as the long-term effects of neonatal ethanol exposure on parvalbumin-interneuron mediated neurotransmission in adolescent mice. Mice were exposed to ethanol using vapor inhalation chambers. In postnatal day (P) 7 mouse pups, ethanol unexpectedly failed to potentiate GABA_A_ receptor-mediated synaptic transmission. Binge-like ethanol exposure of P7 mice expressing channel rhodopsin in parvalbumin-positive interneurons enhanced the peak amplitudes, asynchronous activity and total charge, while decreasing the rise-times of optically-evoked GABA_A_ receptor-mediated inhibitory postsynaptic currents in adolescent animals. These effects could partially explain the learning and memory deficits that have been documented in adolescent and young adult mice exposed to ethanol during the third trimester-equivalent developmental period.

## INTRODUCTION

Exposure to ethanol during fetal development causes a spectrum of deficits, including growth retardation, craniofacial anomalies, and CNS alterations. Ethanol affects multiple developmental processes in the fetal brain, leading to long-lasting neurobehavioral alterations that can have a negative impact on the quality of life. Learning, memory, planning, judgement, attention, fine motor coordination, social interactions, sleep, and emotional control are among the myriad brain functions that can be disrupted by prenatal ethanol exposure. Studies indicate that dysfunction of GABAergic interneurons (INs) likely contributes to these neurobehavioral deficits. Moore et al^1,2^ reported a reduction in the number of parvalbumin-expressing GABAergic INs (PV-INs) in the medial septum and anterior cingulate cortex of adult rats that were exposed to ethanol (peak blood ethanol concentration (BEC) = 160 mg/dl) via liquid diet during gestational days 0 to 21 (i.e., equivalent to the first and second trimesters of human pregnancy). Bailey et al^3^ found a reduction in the number of glutamic acid decarboxylase immunopositive cells in layers II/III of the somatosensory cortex of adult guinea pigs that were exposed to ethanol (oral administration; peak BEC = 328 mg/dl) during the equivalent of all trimesters of human gestation. Miller^4^ observed a reduction in the number of GABA-positive neurons in all layers (except for layer V) of the somatosensory and motor cortices of adolescent macaques that were exposed to ethanol (intragastric intubation; peak BEC = 230 mg/dl) during the first six weeks of gestation or throughout gestation (24 weeks). Cuzon et al^5^ demonstrated that exposure of mice to a low dose of ethanol (liquid diet; peak BEC = 25 mg/dl) during the first 14.5 days of gestation induces premature GABAergic IN tangential migration into the cortical anlage (detected at embryonic day 14.5), an effect that is mediated by increased ambient GABA levels and GABA sensitivity of migrating INs. Skorput et al^6^ showed that 3-day binge-like ethanol exposure during embryonic days 13.5 and 16.5 (liquid diet; peak BEC = 80 mg/dl) increases the density of median ganglionic eminence-derived INs in 16.5-day-old embryos. This effect lingered until young adulthood, as evidenced by both an increase in the number PV-INs in layer V of the medial prefrontal cortex, as well as potentiation of GABA_A_ receptor-mediated synaptic transmission at pyramidal neurons. The mechanism responsible for this effect of ethanol involves potentiation of the depolarizing action of GABA_A_ receptors in migrating cells and increased neurogenesis in the medial ganglionic eminence^7,8^. Larsen et al^9^ exposed differentiating human pluripotent stem cells to ethanol (50 mM for 50 days to model exposure during the first trimester of human gestation) and found a reduction in transcripts related to GABAergic IN specification (i.e., *GSX2, DLX1-6, SST*, and *NPY*) without effects on IN number. Collectively, these studies indicate that ethanol exposure during the equivalent to the first and second trimesters of human pregnancy causes dose- and region-specific effects on IN proliferation, differentiation, migration, and/or survival.

Ethanol exposure during the equivalent to the last trimester of human pregnancy has also been shown to have deleterious effects on INs. Exposure of rats to ethanol vapor (peak BEC = 206 mg/dl) between postnatal day (P) 2 and P6 caused an increase in the number of calretinin positive INs and a reduction in calbindin positive INs (with no change in PV-INs) in the primary motor and somatosensory cortex at P60^10^; this group of investigators also found a reduction in the dendritic tree of PV-INs in the striatum at P60^11^. Also using an ethanol vapor chamber paradigm (E12-19 and P2-9; peak BEC = 330 mg/dl at P7-8), Nirgudkar et al^12^ found reductions in cerebellar IN numbers at P16. Bird et al^13^ demonstrated that ethanol vapor exposure during P2-9 (peak BEC = 221 mg/dl) reduces IN numbers in the adult mouse hippocampus; this study also found that a single vapor chamber exposure at P7 (peak BEC = 297 mg/dl) increases the number of INs that express activated caspase-3, suggesting that they are programmed to undergo apoptotic neurodegeneration. Ethanol administration to P7 mice (subcutaneous injection; peak BEC near 500 mg/dl) has been shown to reduce the numbers of PV-INs in the frontal cortex at P82^14^, as well as in the hippocampal formation (at P14 and P90-100) and the pyriform cortex (at P100)^15,16^ The same subcutaneous P7 ethanol administration paradigm has been demonstrated to decrease the numbers of PV and calretinin positive INs in the neocortex of adult mice^17^. It can be concluded from these results that INs are particularly sensitive targets of the effects of ethanol exposure during the third trimester-equivalent of human pregnancy.

Studies with humans and animals have demonstrated that fetal ethanol exposure causes learning and memory deficits and that these could be, in part, a consequence of alterations in hippocampal function (reviewed in^18,19^). Comparatively, little is known about the contribution to these deficits of alterations in the function of other cortical regions that interconnect with the hippocampus. One of such regions is the retrosplenial cortex (RSC), which plays a central role in navigation, spatial learning, memory, and contextual fear conditioning^20,21^. Studies with mice have demonstrated that heavy binge-like exposure to ethanol at P7 triggers apoptosis of pyramidal neurons and PV-INs in the RSC, an effect that could, in part, underlie the deficits in contextual fear conditioning and navigation in the Morris water maze caused by this ethanol exposure paradigm^22–27^. Consistent with these studies, we recently reported that P7 ethanol vapor administration (peak BEC = 400 mg/dl) triggers apoptotic neurodegeneration of INs in the murine RSC^28^. In the same study, we also found that acute bath application of ethanol to brain slices from P6-8 mice decreased the amplitude of both NMDA and non-NMDA glutamatergic excitatory postsynaptic currents; however, only inhibition of NMDA receptors affected synaptic excitability in RSC neurons. These results support the hypothesis that inhibition of NMDA receptors mediates the apoptogenic effects of third-trimester ethanol exposure^22,29–31^. Here, we tested whether this ethanol exposure paradigm causes either acute or long-lasting alterations in GABA_A_ receptor-mediated neurotransmission in the RSC. Specifically, to determine the acute effects of ethanol on GABA_A_ receptors, we first bath applied ethanol to tissue slices from P6-8 pups and measured the effect of ethanol on evoked GABA_A_ receptor-mediated postsynaptic currents. Next, to determine long-term effects of ethanol, we used optogenetic techniques in acute brain slices to investigate its impact on GABAergic transmission at PV-IN-to-pyramidal neurons synapses in young-adult mice.

## METHODS

All procedures involving animals were approved by the Institutional Animal Care and Use Committee of the University of New Mexico Health Sciences Center and adhered to the U.S. Public health Service policy on humane care and use of laboratory animals. All chemicals were purchased from Sigma-Aldrich (St. Louis, MO) unless otherwise indicated.

### Experiment 1

#### Animals

Transgenic mice expressing Venus yellow fluorescent protein in GABAergic and glycinergic INs expressing the vesicular GABA transporter (VGAT) were generously provided by Dr. Yuchio Yanagawa (Gunma University Graduate School of Medicine, Maebashi, Japan)^32^. Mice were housed at 22°C on a reverse 12-h light/dark cycle (lights on at 8 p.m.) with *ad libitum* access to standard chow and water. Both male and female pups P6-P8) were used for Experiment 1 (Figure 1a).

**Figure 1:**
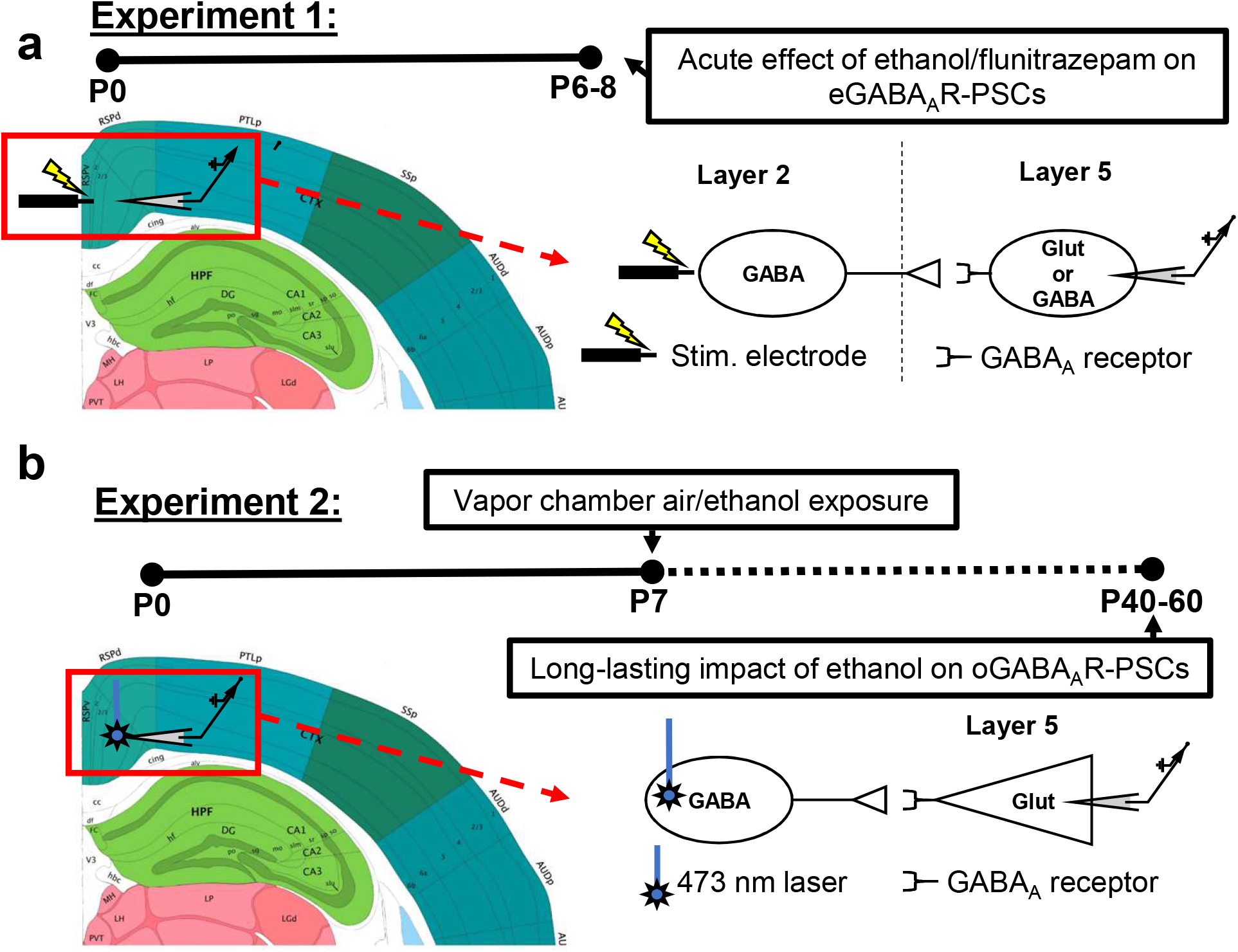
Overview of electrode placement and timeline for the two experiments performed. a) Electrode placement and timeline for Experiment 1. The acute effects of ethanol and flunitrazepam on GABA_A_ receptor mediated PSCs were measured in layer V pyramidal neurons and INs of the RSC. To electrically evoke GABA_A_-PSCs, a bipolar stimulating electrode was placed on the border of layers I/II. b) Electrode placement, laser stimulation location, and timeline for Experiment 2. The long-term effects of P7 ethanol vapor chamber exposure were measured at P40-60. PV-INs expressing channel rhodopsin-2/tdTomato were stimulated with a 473 nm laser using a 40X objective lens in layer V of the RSC. Optically evoked IPSCs were recorded in layer V pyramidal neurons. Atlas images were adapted from the Allen Brain Coronal Atlas https://mouse.brain-map.org/static/atlas^81^. Image credit: Allen Institute.

#### Slice electrophysiology

Slice electrophysiology experiments were performed as described previously^28^. Briefly, P6-P8 pups were anaesthetized with isoflurane (Piramal Critical Care, Bethelhem, PA) and rapidly decapitated. Brains were quickly removed and submerged for 3 min in a protective sucrose cutting solution. Ventral RSC-containing (equivalent to bregma −1.31 to −2.53 in an adult mouse^33^) coronal slices (300 μm) were prepared using a vibrating slicer (VT 1000S, Leica Microsystems, Bannockburn, IL). Brain slices were then placed in a holding solution for 40 min at 32-34°C, followed by storage in a holding solution at room temperature (~24°C) for 30 min before recordings began. Recordings were acquired at 10 kHz and filtered at 2 kHz. Recordings were collected and analyzed using Clampex and Clampfit software (version 10 or 11; Molecular Devices, San Jose, CA), respectively.

Whole-cell patch-clamp recordings of both spontaneous GABA_A_ receptor-mediated postsynaptic currents (GABA_A_-sPSCs) and electrically-evoked GABA_A_ receptor-mediated postsynaptic currents (GABA_A_-ePSCs) were performed using an internal solution composed of (in mM): 140 Cs-methanesulfonate, 0.5 EGTA, 15 HEPES, 2 tetramethylammonium chloride, 2 Mg-ATP, 0.3 Na-GTP, 10 phosphocreatine disodium salt, and 4 QX-314-Cl (Hello Bio, Princeton, NJ) pH 7.25 (adjusted with CsOH) and 305 mOsm. Neurons were equilibrated with the internal solution for 5 min prior to recording. After the equilibration period, the holding potential was gradually increased from −70 mV to 0 mV in order to see GABA_A_-PSCs. Recordings were obtained from pyramidal neurons (Venus negative) and INs (Venus positive) in layer V of the RSC, as well as hippocampal CA1 pyramidal neurons. Evoked PSCs were triggered with a concentric bipolar stimulating electrode (Catalog #CBAEC75, FHC, Bowdoin, ME) that was placed in layer II (or in *stratum radiatum* for CA1 pyramidal neurons) approximately 100 μm away from the cell being recorded from using a MP-225 micromanipulator (Sutter Instruments, Novato, CA) (Figure 1a). Submaximal ePSCs were evoked with a Master-8 pulse stimulator (AMPI, Jerusalem, Israel) connected to an ISO-Flex stimulus isolator (AMPI) every 30 s using 75 μs square wave pulses. Recording of both sPSCs and ePSCs consisted of four phases: 10 min baseline, 10 min drug application, 10 min washout, and finally application of gabazine (25 μM; Hello Bio) to verify that PSCs were indeed GABA_A_ receptor mediated. Kynurenic acid (3 mM) was applied during the equilibration and all recording phases to block excitatory glutamatergic activity. During the drug application phase, either 90 mM ethanol or 1 μM flunitrazepam (Research Biochemicals International, Natick, MA) were applied to the slice.

For sPSC recordings, the last 5 minutes of the baseline, 90 mM ethanol application, and washout phases were analyzed using the Mini Analysis Program (Synaptosoft, Decatur, GA). For ePSC recordings, an input/output curve was initially measured and the stimulation intensity was subsequently adjusted so that the ePSC amplitude was 30-40% of the maximum amplitude. PSCs were evoked at 0.033 Hz. Amplitudes were normalized to the average amplitude of the sPSCs from the entire baseline phase. For ePSCs, normalized amplitude and decay constant tau were measured during the last 5 minutes of each recording phase for both 90 mM ethanol and 1 μM flunitrazepam application experiments. Tau was calculated by averaging the ePSC waveforms from the last 5 minutes of each phase and fitting the resulting trace with a single exponential decay function. In CA1 pyramidal neurons, ethanol had a more transient effect; therefore, its effects on amplitudes and decay taus were calculated from the last 5 minutes of the baseline phase and the first 5 minutes of the 90 mM ethanol application and washout phases.

### Experiment 2

#### Animals

Mice expressing the channel rhodopsin-2 (ChR2) variant ChR2 H134R fused to tdTomato in PV-INs were generated by crossing female heterozygous B6.129P2-Pvalb^tm1(cre)Arbr^/J (B6 PV^cre^, Jackson Laboratory, Bar Harbor, ME; stock number 017320) and male heterozygous B6.Cg-Gt(ROSA)26Sor^tm27.1(CAG-COP4*H134R/tdTomato)Hze^/J (Ai27D, Jackson Laboratory; stock number 012567) mice. After weaning, the offspring (hereafter referred to as B6 PV^cre^-Ai27D mice) were ear-tagged and tail snips were collected under isoflurane anesthesia. Genotyping was performed by Transnetyx (Cordova, TN). Male and female mice aged between P40 and P60 were used for all subsequent experiments (Figure 1b).

#### Ethanol vapor chamber exposure

Pups and dams were exposed to either air or vaporized ethanol (95%, Koptec, King of Prussia, PA) for 4 h (approximately 10 a.m. to 2 p.m.) at P7 in custom-built ethanol vapor chambers^34^. Vapor inhalation chambers have been previously used in multiple laboratories, including our own, to characterize the effects of ethanol exposure on nervous system development^35–43^. Ethanol vapor concentrations were determined using a breathalyzer (Intoximeters, St. Louis, MO) and were between 8-9 g/dl. This exposure paradigm produces peak BECs in pups near 80 mM and triggers apoptotic neurodegeneration in several brain regions, including the RSC^13,28^.

#### Immunohistochemistry

Tissue sections were stained with an anti-PV antibody to verify both the specificity and penetrance of ChR2-H134R/tdTomato expression. Immunohistochemistry (IHC) experiments were performed as described previously^13^. Male and female B6 PV^cre^-Ai27D mice aged P40-60, that were exposed to either air or vaporized ethanol at P7, were deeply anaesthetized with ketamine (250 mg/kg intraperitoneally), the diaphragm was sectioned, and brains were transcardially perfused with 4% paraformaldehyde (PFA, 4% w/v in phosphate buffered saline (PBS), pH 7.4). Brains were extracted and stored in 4% PFA for 48 h, before being cryoprotected for 48 h in 30% sucrose (w/v in PBS). Brains were frozen at −80°C until sectioning. Parasagittal sections (50 μm) were prepared using a cryostat (Microm 505E, Waldorf, Germany). Floating sections were stored at −20°C in a cryoprotectant solution (0.05 M phosphate buffer pH 7.4, 25% glycerol and 25% ethylene glycol). Five randomly selected parasagittal sections (from both hemispheres) containing the ventral RSC were incubated for 2 h in 1% bovine serum albumin, 0.2% Triton X-100, and 5% donkey serum (Jackson ImmunoResearch, West Grove, PA) in PBS (pH 7.4). The sections were next incubated with a 1:5,000 dilution of anti-PV monoclonal antibody (catalog # 235, Swant Inc., Switzerland) at 4°C. Sections were then incubated in a secondary antibody solution containing a 1:1,000 dilution of donkey anti-mouse IgG Alexa Fluor™ 488 antibody (catalog # A21202, Thermo-Fisher, Waltham, MA,) for 2 h at room temperature (approximately 23°C), and then for 20 min in 600 nM 4’6-diamidino-2-phenylindole hydrochloride (DAPI). Sections were rinsed with PBS and mounted on Superfrost Plus microscope slides (VWR, Radnor, PA) using Fluromount G mounting media (Southern Biotech, Birmingham, AL) and covered with glass coverslips (VWR).

Sections were imaged using a Nuance spectral imaging system (PerkinElmer, Hopkinton, MA) on a Nikon TE-200 U inverted fluorescence microscope (Nikon Instruments Inc., Melville, NY) as described previously^28^. Images were acquired with a 40X objective (Plan-NEOFLUAR 40X/1.3 oil, Zeiss, White Plains, NY). The number of Alexa-Fluor 488 stained PV positive cells (525 nm emission maximum), the number of cells expressing tdTomato (581 nm emission maximum), and the number of cells positive for both for Alexa-Fluor 488 and tdTomato were exhaustively counted in Layer V of the ventral RSC in each section by an experimenter blinded to the experimental conditions. For each animal, the percentage of PV positive cells colocalized with tdTomato were ascertained as a measure of transgene penetrance (i.e., number of PV+ cells colocalized with tdTomato divided by the number of PV+ cells), and the percentage of cells expressing tdTomato that did not express PV was determined as a measure of transgene specificity (i.e, number of cells expressing only tdTomato without PV divided by the number of PV+ cells colocalized with tdTomato). Cells were counted using the point selection tool in Fiji (NIH Image J software^44^).

#### Optogenetic slice electrophysiology

Electrophysiology experiments were performed using the same methods as described in Experiment 1 with the modifications detailed below. Slices were prepared from male and female B6 PV^cre^-Ai27D mice aged P40-P60 using the protective cutting/recovery methodology of Ting et al^45^. Mice were deeply anaesthetized with ketamine (250 mg/kg intraperitoneally) and transcardially perfused with 25 mL of a protective N-methyl-D-glucamine (NMDG)-containing aCSF at 4°C composed of (in mM): 92 NMDG, 2.5 KCl, 1.25 NaH_2_PO_4_, 30 NaHCO_3_, 20 HEPES, 25 glucose, 2 thiourea, 3 sodium pyruvate, 5 ascorbic acid, 10 MgSO_4_, and 0.5 CaCl_2_ saturated with 95% O_2_/5% CO_2_ (pH 7.3-7.4 with HCl; 300-310 mOsm). Brains were rapidly removed and immersed for 1 min in the same NMDG-containing aCSF. Coronal brain slices (300 μM) were prepared using a vibrating slicer (described above under Experiment 1) in NMDG aCSF at 4°C. Once all slices containing the ventral RSC (bregma −1.31 to − 2.53^33^) were prepared, they were transferred to warm NMDG aCSF holding solution at 32-34°C. Over the course of 25 min, the NaCl concentration of the warm holding solution was gradually increased to 52 mM by adding increasing amounts of NMDG-aCSF containing 2M NaCl (250 μl at 0 min, 250 μl at 5 min, 500 μl at 10 min, 1 ml at 15 min, and 2 ml at 20 min). Slices were then allowed to recover in holding solution at room temperature for 1 h (as described for Experiment 1 above).

Following recovery, slices were transferred to the recording chamber and aCSF containing 3 mM kynurenic acid was applied at a rate of 2 ml/min. Recording electrodes were filled with the same Cs-methanesulfonate internal solution used in Experiment 1. Optically-evoked inhibitory PSCs (oIPSCs) were generated in layer V pyramidal neurons using a 473 nm laser (IKE-473-100-OP) connected to a power supply (IKE PS-300) (IkeCool Corporation, Anaheim, CA) (Figure 1b). Laser light was delivered through the 40X objective lens using an IS-OGP optogenetics laser positioner (Siskyou, Grants Pass, OR). Laser output power was 20 mW. Cellular capacitance and membrane resistance were measured immediately before the oIPSC recordings began. Three oIPSCs at 20-s intervals were generated at each of the following laser pulse durations: 0.5 ms, 1 ms, 2 ms, 4 ms, and 8 ms. The oIPSC was then blocked with gabazine (25 μM) to confirm that the oIPSC was a GABA mediated current. Data from any cell with an oIPSC that was not blocked with gabazine were discarded. The average oIPSC at each laser pulse duration for every cell was analyzed for peak amplitude, current density, GABAergic total charge (area under the curve, pA X ms), half-width at half-maximal amplitude, and rise time using Clampfit (Molecular Devices). In addition, the number of individual oIPSC peaks for each evoked current was counted for 100 ms after the onset of the laser stimulation, and the results were averaged together within each laser pulse duration for each cell. To measure oIPSC paired pulse ratios (PPRs), we evoked 10 pairs of oIPSCs at 30 s intervals, with 50 ms between paired laser pulses and 1 ms laser pulse durations. The ratio of the amplitude or the total charge of the second peak divided by the first peak (P2/P1) was measured.

#### Statistics

Statistical analyses were performed using Prism version 8.4.2 (GraphPad Software, San Diego, CA) and SPSS version 26 (IBM, Armonk, NY). Precise p-values are reported as recommended by a recent article^46^. Data from Experiment 1 were analyzed using repeated measures ANOVA with cell-type (pyramidal or IN) as the fixed factor and the three phases (baseline, drug application, washout) as the repeated measure. ANOVA residuals were tested for normality using a Shapiro-Wilkes test, and any data that failed this test (p < 0.05) was analyzed using a non-parametric Friedman’s ANOVA test. Since Experiment 1 examined the acute effect of ethanol or flunitrazepam on PSCs, the unit of determination was a single cell. For ePSCs, the dependent measure was either evoked current amplitude or decay (tau). For sPSCs, the dependent measures were frequency, amplitude, rise time, and decay (tau). Main effect of cell-type for non-parametric data was analyzed using a Mann-Whitney U test. F-ratios and p-values are reported for cell-type, phase, and cell-type by phase interactions for all dependent variables. Greenhouse-Geisser corrected F-ratios and p-values are reported for repeated measures that failed (p < 0.05) Mauchley’s test of sphericity. Any cell-type X phase interactions were further explored by performing a multiple comparison test within cell type, comparing the baseline, drug-application and washout in a pairwise manner. Bonferroni corrected p-values are reported for all multiple comparison tests. Effect sizes for Experiment 1 are reported as follows: partial eta squared (η_p_^2^) for ANOVAs, Kendall’s *W* for Friedman’s ANOVA, Hedge’s *g* for pairwise multiple comparison post hoc tests, and *r* for Friedman’s ANOVA post hoc multiple comparisons and Mann-Whitney U tests.

IHC data from Experiment 2 was analyzed using an unpaired t-test for parametric data or a Mann-Whitney U test for non-parametric data. Effect sizes for these tests are reported as Hedges’ *g* or *r*, respectively. For optogenetic electrophysiology experiments from Experiment 2, because multiple cells were recorded from several animals from several litters, we used a linear mixed-model (LMM) approach to data analysis in SPSS adapted from procedures described by West et al^47^. This approach allows us to determine if including random litter effects significantly improves the LMM being used for a particular dependent variable^48^. Detailed statistics from LMM analyses appear in Supplemental Table 2. Before LMM analyses were performed, outliers for each dependent variable were removed using a 1% ROUT test in GraphPad. The fixed factors were sex, P7 vapor chamber exposure condition, and laser pulse duration as a repeated measure. Models were built stepwise for each dependent variable according to the following procedure. First, a LMM was fit that includes the random effect associated with the intercept for each litter, using homogenous residual error variances between treatment groups, and an unstructured covariance structure for the repeated measures residuals. If the LMM failed to achieve convergence using an unstructured covariance structure for repeated measures residuals, a compound symmetry covariance structure was used instead. Then, a LMM was fit without the random effect included. The −2-log restricted maximum likelihood value for each model fit was then used to perform a likelihood ratio chi-square test. If the p-value for this test was < 0.05, this indicated that random effects significantly improved the model fit and were therefore included in subsequent models. The resulting LMM was then fit using heterogenous residual error variances for treatment groups, and another likelihood ratio chi-square test was performed. If the p-value for this test was < 0.05, including heterogenous residual error variances significantly improved the model and subsequent LMMs included heterogenous residual error variances. F-ratios (containing Satterthwaite approximated degrees of freedom) and p-values for Type III F-tests for treatment, sex, laser pulse duration, and interactions between fixed factors from the final LMM are reported in the results section for each dependent variable. Effect sizes for main effects of exposure and sex for LMMs are reported as Hedges’ *g* and not partial eta squared because SPSS currently does not provide a sum of squares output for these tests. Some skewness and kurtosis in data distribution can be tolerated using LMMs as they are robust against violations of assumptions of normality, as normality of residuals does not affect parameter estimates in multilevel models^49^. However, because residuals from many of the LMMs violated assumptions of normality (Shapiro-Wilkes p-value of < 0.05), we also present non-parametric Mann-Whitney U tests for main effects of vapor chamber exposure condition and sex in Supplemental Table 2 for any LMM that did not pass the Shapiro-Wilkes test. Exposure by laser pulse duration interactions with a p-value of < 0.05 were subsequently analyzed with non-parametric Mann-Whitney U tests examining the effect of vapor chamber exposure condition within each laser pulse duration. P-values reported for these tests are Bonferroni corrected. Detailed statistics for post hoc tests examining exposure effects within each laser pulse duration appear in Supplemental Table 3. All data presented are mean ± standard error of the mean (SEM).

## RESULTS

### Experiment 1: Effects of acute ethanol exposure on GABA_A_ receptor-mediated PSCs in the RSC

In our previous work^28^, we demonstrated that bath application of 90 mM ethanol to acutely prepared brain slices from P6-P8 pups inhibited NMDA and non-NMDA receptor-mediated excitatory postsynaptic currents in both pyramidal neurons and INs from layer V of the RSC. This concentration is above the threshold of BECs required for the induction of apoptotic neurodegeneration in postnatal rodents^13,22,28^. In this work, we sought to determine the effects of bath-applied ethanol on GABA_A_-ePSCs in pyramidal neurons and INs in the same region. We hypothesized that ethanol acutely potentiates GABA_A_ receptors, which could contribute to apoptosis caused by excessive inhibition of neuronal activity. We applied 90 mM ethanol to acutely prepared brain slices from male and female mice at P6-P8. We elicited submaximal GABA_A_-ePSCs in layer V pyramidal neurons and INs using a bipolar stimulating electrode placed in layer II (Figure 1a). Average traces from pyramidal neurons and INs obtained during the baseline and 90 mM ethanol exposure phases appear in Figures 2a and 2b, respectively. Contrary to our expectations, acute bath application of 90 mM ethanol failed to potentiate GABA_A_-ePSC amplitudes in either pyramidal neurons or INs in layer V of the RSC (Figure 2c). Using a repeated-measures two-way ANOVA, we observed an effect of exposure phase, with amplitudes decreasing over the course of the experiment instead of increasing during the 90 mM ethanol exposure phase (phase F(1.421’22.742) = 3.934, p = 0.047, η_p_^2^ = 0.197; male pyramidal n = 4 cells from 4 animals from 2 litters, female pyramidal n = 6 cells from 5 animals from 4 litters; male IN n = 4 cells from 4 animals from 2 litters; female IN n = 4 cells from 4 animals from 3 litters). There was an effect of cell type examined, as overall ePSC amplitudes were larger in pyramidal neurons (501.3 ± 118.3 pA) than in INs (176.4 ± 58.7 pA) (cell type F(1,16) = 5.225, p = 0.036, η_p_^2^ = 0.246), but no interaction between exposure phase and cell type (interaction F(1.421, 22.742) = 0.369, p = 0.62, η_p_^2^ = 0.023).

**Figure 2:**
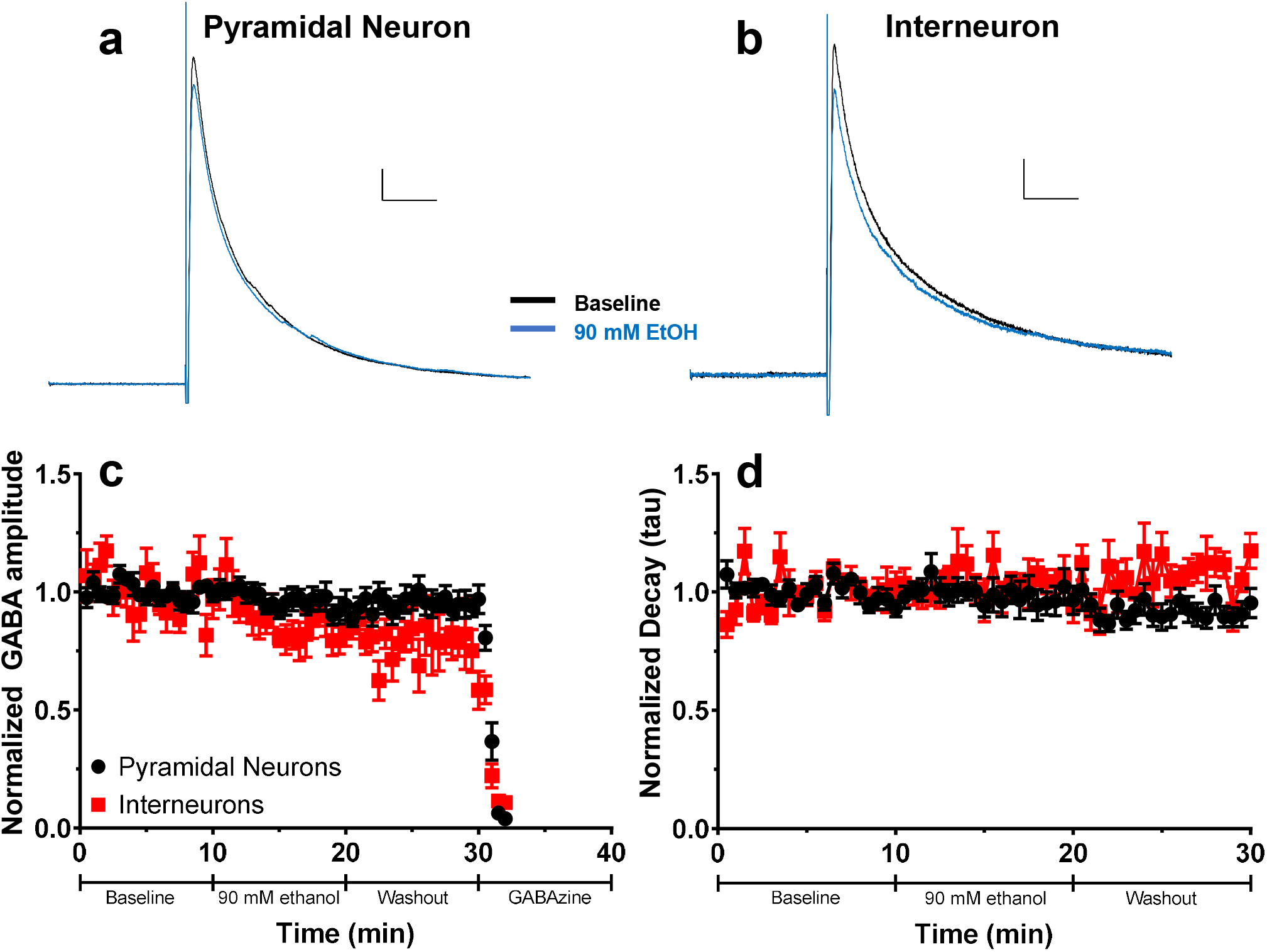
Effect of acute ethanol application on evoked GABA_A_ receptor-mediated postsynaptic current (GABA_A_-ePSC) amplitude and decay in layer V pyramidal neurons and INs. a) Average ePSC traces from layer V pyramidal neurons during the baseline (black trace) and acute 90 mM ethanol application (blue trace) phases. Scale bars = 40 ms, 50 pA. b) Average ePSC traces from layer V INs during the baseline and acute 90 mM ethanol application phases. Scale bars = 40 ms, 20 pA. c) Normalized GABA_A_-ePSC amplitudes for both pyramidal neurons (black circles) and INs (red squares) in layer V during the baseline, 90 mM ethanol application, washout, and gabazine (25 μM) application phases. d) Normalized decay constants (tau) for both pyramidal neurons and INs during the baseline, 90 mM ethanol application and washout phases. Data are presented as mean ± SEM.

Although ethanol does not potentiate GABA_A_-ePSCs in RSC neurons by increasing current amplitude, it may increase the decay time of the current, allowing for more total current flow into the postsynaptic cell. Therefore, GABA_A_-ePSCs were fit with a single exponential decay curve, and the decay constant tau was measured. Bath application of 90 mM ethanol failed to affect the decay constant of GABA_A_-ePSCs in either cell type examined (Figure 2d). Ethanol application and washout did not affect GABA_A_-ePSC decay (Friedman’s ANOVA χ^2^(2) = 2.111, p = 0.35, *W* = 0.059). There was no effect of cell type examined (pyramidal neuron tau 30.96 ± 3.25; IN tau 41.53 ± 6.97) (Mann-Whitney U (n1 = 10, n2 = 8) = 25, p = 0.20, *r* = 0.314).

As a positive control, we also recorded GABA_A_-ePSCs from hippocampal CA1 pyramidal neurons, as it has been shown that acute ethanol application can potentiate these currents^50,51^. In these cells, 90 mM ethanol application potentiated GABA_A_-ePSCs for a relatively short duration. For this reason, we compared ePSC amplitudes and decays from the last 5 minutes of the baseline phase to the first 5 minutes of the 90 mM ethanol application and washout phases. Average traces from hippocampal CA1 pyramidal neurons during the baseline and 90 mM application phases appear in Supplemental Figure 1a. Using one-way ANOVA, we observed an effect of phase on CA1 pyramidal neuron ePSC amplitudes (phase F(2,24) = 4.101, p = 0.030, η_p_^2^ = 0.255; Supplemental Figure 1 b). There was no effect of ethanol exposure or washout on ePSC decay tau in CA1 pyramidal neurons (phase F(2,24) = 0.484, p = 0.62, η_p_^2^ = 0.039; Supplemental Figure 1c). These results demonstrate that 90 mM ethanol application has the ability to potentiate GABA_A_-ePSCs in neurons outside of layer V of the RSC.

After observing that 90 mM ethanol application did not potentiate GABA_A_-ePSCs in either pyramidal neurons or INs in layer V of the RSC, we confirmed that it was possible to potentiate GABA_A_-ePSCs in these developing neurons. To this end, we performed the same experiment described above using the allosteric potentiator of GABA_A_ receptors, flunitrazepam (1 μM). Flunitrazepam is a benzodiazepine that has been demonstrated to increase decay time of GABA_A_-PSCs (Bouairi et al., 2006). Average traces during the baseline and 1 μM flunitrazepam application phases for pyramidal neurons and INs appear in Figure 3a and Figure 3b, respectively. Application of 1 μM flunitrazepam did not increase the amplitude of the ePSCs from either cell type (phase F(1.235,32.117) = 2.000, p = 0.16, η_p_^2^ = 0.071; interaction F(1.235,32.117) = 0.376, p = 0.59, η_p_^2^ = 0.014; male pyramidal n = 9 cells from 6 animals from 4 litters; female pyramidal n = 5 cells from 3 animals from 3 litters; male IN n = 10 cells from 6 animals from 4 litters; female IN n = 4 cells from 2 animals from 2 litters) (Figure 3c) There was an effect of cell type on ePSC amplitudes (cell type F(1,26) = 11.562, p = 0.002, η_p_^2^ = 0.308), as overall ePSC amplitudes were larger in pyramidal neurons (510.0 ± 67.9 pA) than in INs (240.1 ± 41.1 pA).

**Figure 3:**
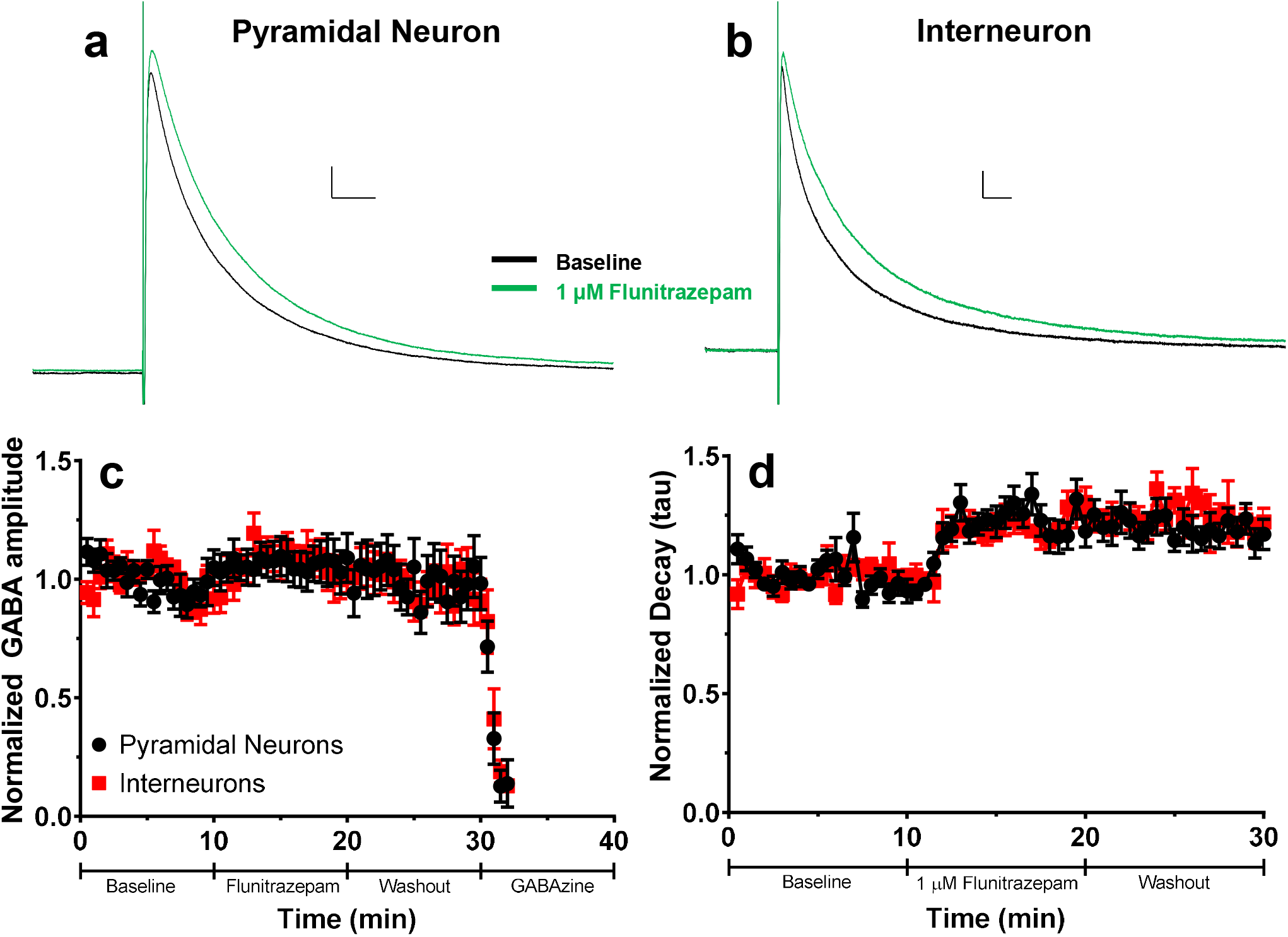
Effect of acute flunitrazepam application on evoked GABA_A_ receptor-mediated postsynaptic current (GABA_A_-ePSC) amplitude and decay in layer V pyramidal neurons and INs. a) Average ePSC traces from layer V pyramidal neurons during the baseline (black trace) and acute 1 μM flunitrazepam application (green trace) phases. Scale bars = 40 ms, 50 pA. b) Average GABA_A_-ePSC traces from layer V INs during the baseline and acute 1 μM flunitrazepam application phases. Scale bars = 40 ms, 20 pA. c) Normalized GABA_A_-ePSC amplitudes for both pyramidal neurons (black circles) and INs (red squares) in layer V during the baseline, 1 μM flunitrazepam, washout, and gabazine application phases. d) Normalized decay constants (tau) for both pyramidal neurons and INs during the baseline, 1 μM flunitrazepam, and washout phases. Data are presented as mean ± SEM.

Flunitrazepam application, however, increased GABA_A_-ePSC decays in both cell types. The decay constant tau was elevated in both cell types during the 1 μM flunitrazepam application phase, and this potentiation continued into the washout phase (Figure 3d). There was an effect of phase (Friedman’s ANOVA χ^2^(2) = 39.071, p < 0.0001, *W* = 0.698; post hoc baseline vs 1 μM flunitrazepam U(n1 = n2 = 28) z = 5.479, p < 0.0001, *r* = 1.035; post hoc baseline vs wash U(n1 = n2 = 28) z = 5.345, *r* = 1.010) but no differences between cell types (pyramidal neuron tau 70.33 ± 6.28, IN tau 87.92 ± 7.91) (Mann-Whitney U (n1 = n2 = 14) = 62, p = 0.10, *r* = 0.313). These results indicate that GABA_A_ receptors on layer V pyramidal neurons and INs in the RSC of P6-P8 animals can be potentiated by drugs that allosterically modulate GABA_A_ receptor function but that 90 mM ethanol does not potentiate evoked GABA_A_ receptor activity. Therefore, it is unlikely that acute apoptosis resulting from ethanol exposure at this critical developmental age is a result of potentiation of inhibition via GABA_A_ receptors.

To more thoroughly determine that 90 mM ethanol application did not have any potentiating effects on GABA_A_ receptor-mediated synaptic transmission in layer V pyramidal neurons and INs, we recorded GABA_A_-sPSCs during the baseline, 90 mM ethanol bath application, and washout phases. Average GABA_A_-sPSC traces for pyramidal neurons and INs during the baseline, 90 mM ethanol application, and washout phases appear in Figures 4a and 4b, respectively. Representative expanded time scale GABA_A_-sPSC traces for pyramidal neurons and INs during the baseline, 90 mM ethanol application, washout, and 25 μM gabazine application phases appear in Supplemental Figures 2a and 2b, respectively. Representative compressed time scale traces for pyramidal neurons and INs appear in Supplemental Figures 2c and 2d, respectively. We measured the sPSC frequency, amplitude, rise time (10-90%) and single-exponential decay constant tau during each of the phases for the two cell types. Cumulative probability plots and average results from each cell for these measures appear in Figures 4c-j (male pyramidal n = 4 cells from 4 animals from 4 litters; female pyramidal = 4 cells from 4 animals from 3 litters; male IN = 4 cells from 4 animals from 4 litters, female IN = 3 cells from 3 animals from 2 litters). There was no effect of exposure phase on frequency of GABA_A_-sPSCs (phase F(1.382,17.970) = 3.485, p = 0.067, η_p_^2^ = 0.211), and no phase by cell type interaction (F(1.382,17.970) = 0.345, p = 0.63. η_p_^2^ = 0.026) (pyramidal neuron frequency Figure 4c; IN frequency Figure 4d). Pyramidal neuron GABA_A_-sPSCs occurred more frequently than IN GABA_A_-sPSCs (cell type F(1,13) = 9.980, p = 0.0075, η_p_^2^ = 0.434). There was an effect of exposure phase on the amplitude of GABA_A_-sPSCs (phase F(1.336,17.371) = 4.913, p = 0.031, η_p_^2^ = 0.274) and an interaction between exposure phase and cell type (interaction F(1.336,17.371) = 6.108, p = 0.017, η_p_^2^ = 0.320), but no effect of cell type examined (cell type F(1,13) = 0.137, p = 0.72, η_p_^2^ = 0.010) (pyramidal neuron amplitude Figure 4e; IN amplitude Figure 4f). The effect of phase and the interaction between phase and cell type was primarily driven by an increase in IN GABA_A_-sPSC amplitude during the washout phase as compared to the baseline phase, but this effect was not observed using a multiple comparison post hoc test (t(6) = 2.357, p = 0.17, *g* = 0.856). There were no effects on GABA_A_-sPSC rise time (phase F(2,26) = 2.763, p = 0.082, η_p_^2^ = 0.175; cell type F(1,13) = 1.448, p = 0.25, η_p_^2^ = 0.100; interaction F(2,26) = 0.265, p = 0.77, η_p_^2^ = 0.020) (pyramidal neuron rise time Figure 4g; IN rise time Figure 4h). There was also no effect of exposure phase or interaction between exposure phase and cell type on GABA_A_-sPSC decay tau (phase F(2,24) = 0.834, p = 0.45, η_p_^2^ = 0.065; interaction F(2,24) = 0.123, p = 0.89, η_p_^2^ = 0.010) (pyramidal neuron tau Figure 4i; IN tau Figure 4j). INs had a higher tau value than pyramidal neurons, indicating that GABA_A_-sPSCs decayed more slowly in INs than pyramidal cells (cell type F(1,12) = 13.543, p = 0.0031, η_p_^2^ = 0.530).

**Figure 4:**
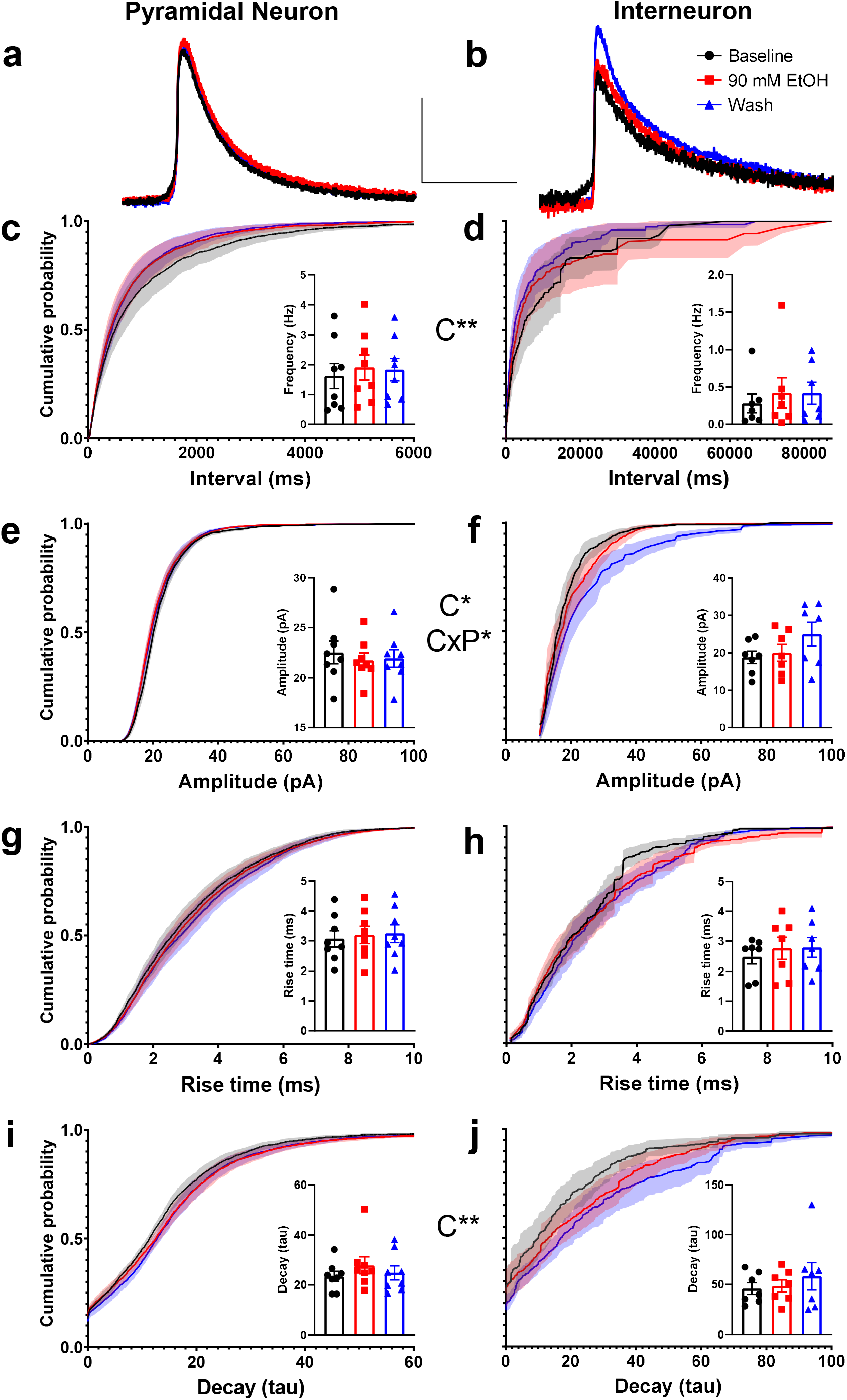
Effect of acute ethanol application on spontaneous GABA_A_ receptor-mediated PSC (GABA_A_-sPSC) characteristics from layer V pyramidal neurons and INs. a) Average GABA_A_-sPSC current traces from pyramidal neurons during baseline (black trace), during 90 mM ethanol bath application (red trace), and during washout (blue trace). b) Average GABA_A_-sPSC current traces from INs during baseline, 90 mM ethanol application, and washout. c-j) Cumulative probability plots and average data from each cell (insets) showing effect of 90 mM ethanol application and washout on c) pyramidal neuron inter-event interval (frequency in inset), d) IN inter-event interval (frequency in inset), e) pyramidal neuron amplitude, f) IN amplitude, g) pyramidal neuron rise time, h) IN rise time, i) pyramidal neuron decay tau, and j) IN decay tau. Baseline = black line/black squares, 90 mM ethanol application = red line/ red circles, washout = blue line/blue triangles. Asterisks (*,**) denote a p-value of p < 0.05 and p < 0.01, respectively. Please see Supplemental Table 1 for exact p-values. Two-Way ANOVA effects: C = cell type effect, P = exposure phase effect, CxP = cell type by exposure phase interaction. Male pyramidal n = 4 cells from 4 animals from 4 litters; female pyramidal = 4 cells from 4 animals from 3 litters; male IN = 4 cells from 4 animals from 4 litters, female IN = 3 cells from 3 animals from 2 litters. Data are presented as mean ± SEM in all cases.

### Experiment 2: Long-term effects of P7 ethanol vapor chamber exposure on oIPSCs in layer V pyramidal neurons in the ventral RSC

Third trimester-equivalent ethanol exposure has been shown to trigger apoptosis and long-term loss of INs, including PV+ INs, in the RSC^25,28^, which could lead to persistent alterations in GABAergic neurotransmission. To examine this possibility, we used a transgenic mouse model (B6 PV^cre^-Ai27D mice) that allowed us to optically stimulate PV+ INs specifically. We assessed whether PV+ IN-mediated oIPSCs in RSC layer V pyramidal neurons of adolescent mice are affected by P7 developmental ethanol exposure. We first determined the penetrance and specificity of transgene expression before starting electrophysiology experiments. We also determined if P7 ethanol exposure altered either of these measures. Male and female mice from both P7 vapor chamber exposure conditions were left undisturbed until P40-P60 and then processed for immunohistochemistry to analyze both PV expression and ChR2/tdTomato expression. Representative IHC images from B6 PV^cre^-Ai27D mice appear in Figure 5a-d. In air exposed animals, mean transgene penetrance was 98.67 ± 1.33% (percentage of PV+ cells expressing ChR2-tdTomato), while in ethanol exposed animals transgene penetrance was 98.82 ± 1.18% (Figure 5e; air male n = 3 animals from 3 litters, air female n = 2 animals from 2 litters; ethanol male n = 3 animals from 3 litters, ethanol female n = 2 animals from 2 litters). Transgene penetrance was not different between vapor-chamber exposure conditions (Mann-Whitney U(n1 = n2 = 5) = 12, p > 0.99, *r* = 0.047). We also measured transgene specificity to see if ChR2-tdTomato was aberrantly expressed in PV-negative cells. In air exposed animals, non-specific transgene expression (% of ChR2-tdTomato positive cells that were not positive for PV) was 3.76 ± 1.58%, while in ethanol exposed animals non-specific transgene expression was 4.81 ± 2.05% (Figure 5f). Transgene specificity was not different between vapor chamber exposure conditions (t(8) = 0.406, p = 0.70, *g* = 0.232).

**Figure 5:**
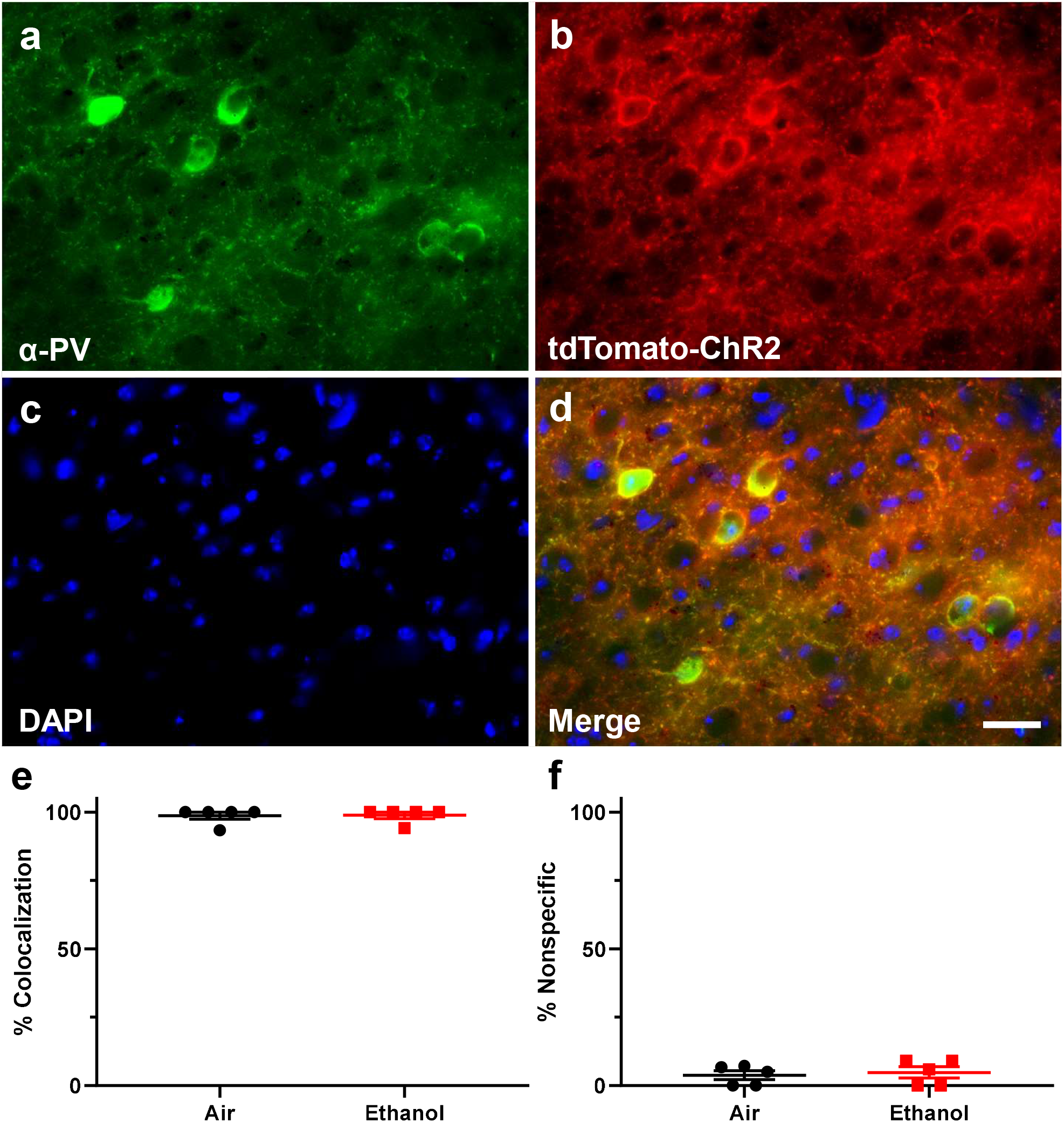
Representative immunohistochemistry (IHC) images from B6 PV^cre^-Ai27D mice and analysis of both penetrance and specificity of transgene expression. a-d) Representative IHC images showing colocalization of PV and ChR2–tdTomato. a) IHC demonstrating PV expression. b) Expression of endogenous ChR2-tdTomato transgene expression. c) DAPI nuclear stain. d) Merged image. Scale bar = 25 μm. e) Penetrance analysis of B6 PV^cre^-Ai27D transgene expression. Penetrance of transgene expression was measured as the number of PV expressing cells colocalized with tdTomato divided by the number of PV expressing cells. f) Specificity analysis of B6 PV^cre^-Ai27D transgene expression. Specificity of transgene expression was measured as the number of cells expressing only tdTomato divided by the number of cells expressing both PV and tdTomato. Air male n = 3 animals from 3 litters, air female n = 2 animals from 2 litters; ethanol male n = 3 animals from 3 litters, ethanol female n = 2 animals from 2 litters. Data are presented as mean ± SEM.

After verifying that B6 PV^cre^-Ai27D mice had acceptable transgene expression, we then examined the effect of P7 ethanol exposure, sex, and laser pulse duration on PV+ IN-mediated oIPSCs in layer V pyramidal neurons in the RSC of both male and female mice at P40-P60 using whole-cell patch-clamp electrophysiology. Average oIPSC traces from each laser pulse duration for mice from air and ethanol vapor chamber exposure conditions appear in Figure 6a and 6b, respectively (female air n = 32 cells from 8 animals from 7 litters; male air n = 46 cells from 9 animals from 8 litters; female ethanol n = 35 cells from 8 animals from 8 litters; male ethanol n = 40 cells from 8 animals from 7 litters). Average oIPSC traces from each laser pulse duration for mice from air and ethanol vapor chamber exposure conditions presented separately by sex appear in Supplemental Figures 3a-d. There was no effect of vapor chamber exposure condition, sex, or interaction between sex and exposure condition on membrane capacitance (p-values > 0.07, Supplemental Figure 3e). There was no effect of vapor chamber exposure condition or interaction between exposure condition and sex on membrane resistance (p-values > 0.39; Supplementary Figure 3f), but females did have a larger membrane resistance than males (F(1,95.267) = 4.199, p = 0.043, *g* = 0.498). Random effect of litter significantly improved the LMM for membrane resistance and was included in the final model; random effect of litter did not significantly improve the LMM for membrane capacitance and was not included in the final model.

**Figure 6:**
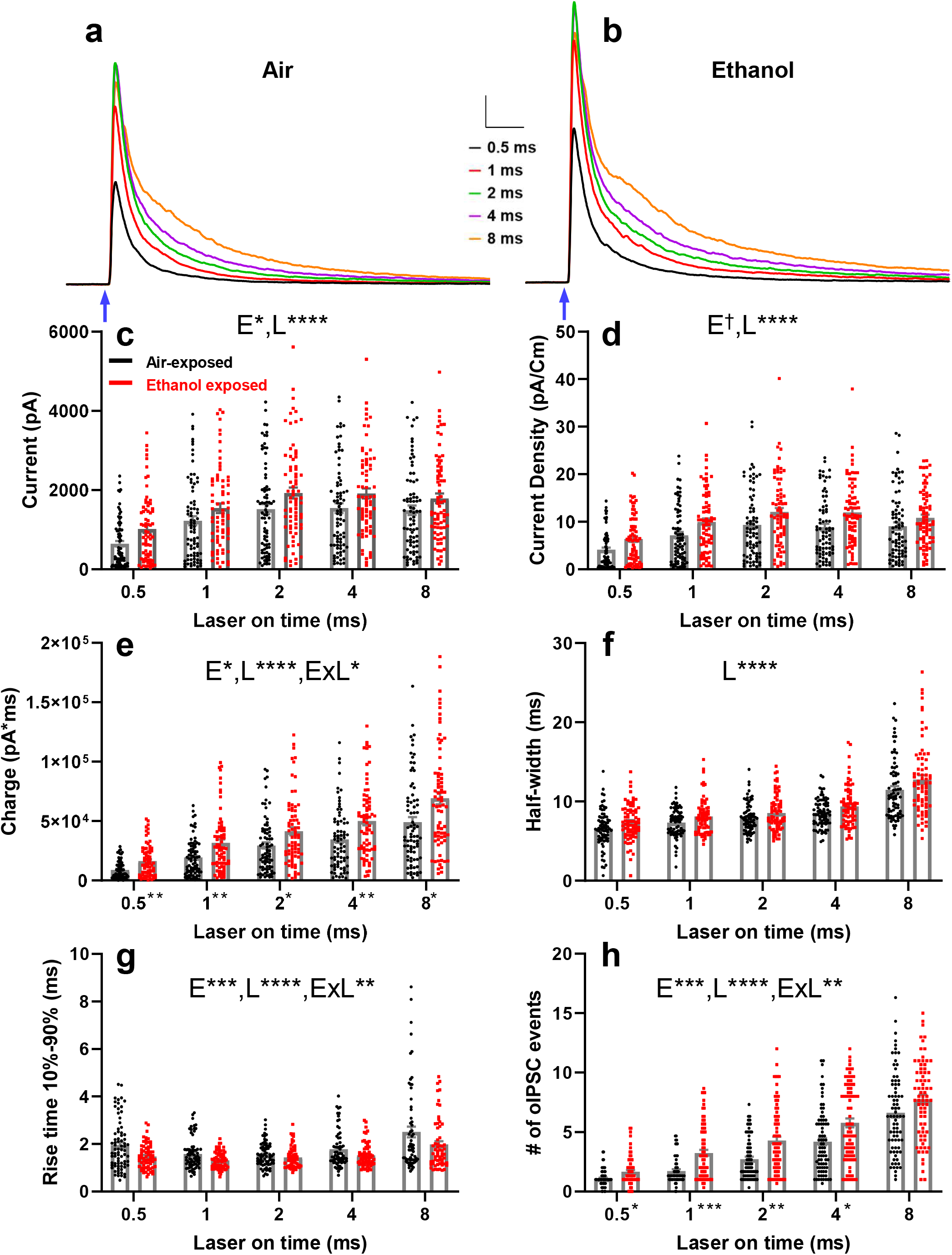
Effect of P7 ethanol exposure on optically-evoked inhibitory postsynaptic currents (oIPSCs) in layer V RSC pyramidal neurons at P40-60. a) Average oIPSC current traces from air-exposed animals using 0.5 ms (black trace), 1 ms (red trace), 2 ms (green trace), 4 ms (purple trace), and 8 ms (orange trace) laser pulse durations. b) Average oIPSC current traces from ethanol-exposed animals using 0.5, 1,2, 4, and 8 ms laser pulse durations. Scale bars = 20 ms, 200 pA. Blue arrows indicate onset of laser pulse. c-h) Collected peak amplitudes (c), current densities (d), GABAergic total charge (e), half widths at half-maximum amplitude (f), rise times (g), and # of oIPSC events (h) for all cells from air exposed (black circles) and ethanol exposed (red squares) animals presented for each laser pulse duration. Data are presented collapsed across sex due to a lack of sex effects from LMM analyses. Dagger (†) denotes a p-value of < 0.06, and asterisks (*,**,***,****) denote p-values of p < 0.05, p < 0.01, p < 0.001, and p < 0.0001, respectively. Please see Supplemental Tables 2 and 3 for exact p-values from LMMs and post hoc tests, respectively. LMM effects: E = exposure effect, L = laser pulse duration effect, ExL = exposure by laser pulse duration interaction. Significance indicators next to x-axis laser pulse duration labels of panel e) and panel h) indicate a Mann-Whitney U post hoc effect of exposure within laser pulse duration. Female air n = 32 cells from 8 animals from 7 litters, male air n = 46 cells from 9 animals from 8 litters; female ethanol n = 35 cells from 8 animals from 8 litters, male ethanol n = 40 cells from 8 animals from 7 litters. Data are presented as mean ± SEM.

Peak amplitude of oIPSCs was increased by vapor-chamber exposure condition (Figure 6c; exposure F(1,146.828) = 5.009, p = 0.027, *g* = 0.328). Peak amplitude was also affected by laser pulse duration (laser F(4,138.866) = 73.641, p < 0.0001) There was no effect of sex, or two- or three-way interactions between sex, exposure, and laser pulse duration on oIPSC amplitude (p-values > 0.08). Random effect of litter did not significantly improve the LMM for oIPSC amplitude and was not included in the final model.

Current density of oIPSCs was also increased in ethanol exposed animals (Figure 6d; exposure F(1,40.564) = 4.044, p = 0.051, *g* = 0.384). Laser pulse duration also affected current density (laser F(4,38.243) = 66.409, p < 0.0001). There was no effect of sex, or two- or three-way interactions between sex exposure, and laser pulse duration on oIPSC current density (p-values > 0.29). Random effect of litter significantly improved the LMM for oIPSC current density and was included in the final model.

The total charge of the oIPSCs was affected by vapor chamber exposure condition. Ethanol exposed animals had a larger oIPSC total charge than air-exposed animals (Figure 6e; exposure F(1,20.848) = 5.907 p = 0.024, *g* = 0.424). Laser pulse duration also affected oIPSC total charge (laser F(4,88.351) = 133.734, p < 0.0001). There was no effect of sex, or interactions between sex and exposure, sex and laser pulse duration, or sex by exposure by laser pulse duration interactions (p-values > 0.23). An exposure by laser pulse duration interaction (F(4,88.351 = 2.849, p = 0.028) was further explored by running Mann-Whitney U tests for exposure effects within each laser pulse duration for total charge. After correcting for multiple comparisons, there was an effect of vapor chamber exposure condition on total charge at the each of the laser pulse durations (all p-values < 0.024, for detailed statistics please see Supplementary Table 3). Random effect of litter for oIPSC total charge significantly improved the LMM and was included in the final model.

The half-width at half-maximal amplitude was not affected by exposure condition (Figure 6f; exposure F(1,26.478) = 2.315, p = 0.14, *g* = 0.270), but was affected by laser pulse duration (laser F(4,40.164 = 51.786, p < 0.0001). There was no effect of sex or interactions between sex, exposure, and laser pulse duration on oIPSC half-width (p-values > 0.32). Random effect of litter significantly improved the LMM for half-width and was included in the final model.

Ethanol vapor chamber exposure decreased the rise time (10-90%) of oIPSCs compared to air-exposed animals (Figure 6g; exposure F(1,110.583) = 11.906, p = 0.0008, *g* = 0.359). Laser pulse duration also affected oIPSC rise time (laser F(4,102.143) = 15.353, p < 0.0001). An exposure by laser pulse duration interaction was further explored using Mann-Whitney U tests for exposure effects within laser pulse duration, but after correcting for multiple comparisons there were no effects (p-values > 0.09; Supplemental Table 3). There was no effect of sex or other interactions between sex, vapor chamber exposure condition, and laser pulse duration for oIPSC rise time (p-values > 0.13). Random effect of litter did not significantly improve the LMM for oIPSC rise time and was not included in the final model.

When analyzing oIPSC data, we noticed an interesting phenomenon. As laser pulse durations increased during the course of an experiment, oIPSC currents contained an increasing number of individual event peaks (Supplemental Figure 4). This effect has been observed previously^52^, and suggests that increases in laser pulse duration promotes asynchronous GABA release following the synchronous release of this transmitter. To analyze this, we counted the number of individual oIPSC events for 100 ms after each laser stimulation and compared experimental groups using a LMM. Ethanol exposure at P7 increased the number of oIPSC events compared to air-exposed animals (Figure 6h; exposure F(1,137.439) = 13.368, p = 0.0004, *g* = 0.387). The number of individual events increased with increases in laser pulse duration (laser F(4,133.965) = 126.460, p < 0.0001). There was no effect of sex, or interactions between sex and exposure, sex and laser pulse duration, or sex by exposure by laser pulse duration interactions (p-values > 0.18). An exposure by laser pulse duration interaction (F(4,133.965) = 3.519, p = 0.0091) was further explored by running Mann-Whitney U tests for exposure effects within each laser pulse duration for number of individual events. After correcting for multiple comparisons, there was an effect of P7 ethanol exposure on number of events for the 0.5, 1,2, and 4 ms laser pulse durations (p-values < 0.009, for detailed statistics please see Supplementary Table 3). Random effect of litter did not significantly improve the LMM for the number of events and was not included in the final model. The presence of the multiple oIPSC peaks made it difficult to determine the effect of ethanol exposure oIPSC decay kinetics, which would have been useful for determining if ethanol exposure causes alterations in postsynaptic GABA_A_ receptor function. The presence of these additional peaks likely contributes to the ethanol-induced increase in oIPSC charge transfer described above, consistent with the role of asynchronous release in prolonging the inhibitory actions of GABA release from INs^53^.

We next analyzed if P7 vapor chamber exposure alters PPRs at PV+ IN-pyramidal neuron synapses. Average PPR current traces from air and ethanol exposed animals show that paired-pulse depression was observed at these synapses, indicating that they have a high basal probability of GABA release (Figure 7a; Air female n = 19 cells from 5 animals from 4 litters, air male n = 22 cells from 6 animals from 5 litters; ethanol female n = 22 cells from 6 animals from 5 litters; ethanol male n = 20 cells from 5 animals from 5 litters). However, ethanol exposure at P7 had no effect on amplitude PPR ratios (Figure 7b; exposure F(1,79) = 0.377, p = 0.54, *g* = 0.159). Female mice had larger PPRs for amplitude than male mice, suggesting that females have a lower basal probability of GABA release (sex F(1,79) = 5.119, p = 0.026, *g* = 0.507). There was no interaction between sex and exposure condition for amplitude PPR (interaction F(1,79) = 0.169, p = 0.68). Similar to amplitude PPR, ethanol exposure had no effect on total charge PPR ratios (Figure 7c, exposure F(1,79) = 1.521, p = 0.22, *g* = 0.291), females had larger charge PPR ratios than male mice (sex F(1,79) = 5.667, p = 0.034, *g* = 0.489), and there was no interaction between sex and exposure condition (interaction F(1,79) = 0.182, p = 0.67). Random effects of litter did not improve the LMM for either amplitude PPR or total charge PPR and were not included in the final model.

**Figure 7:**
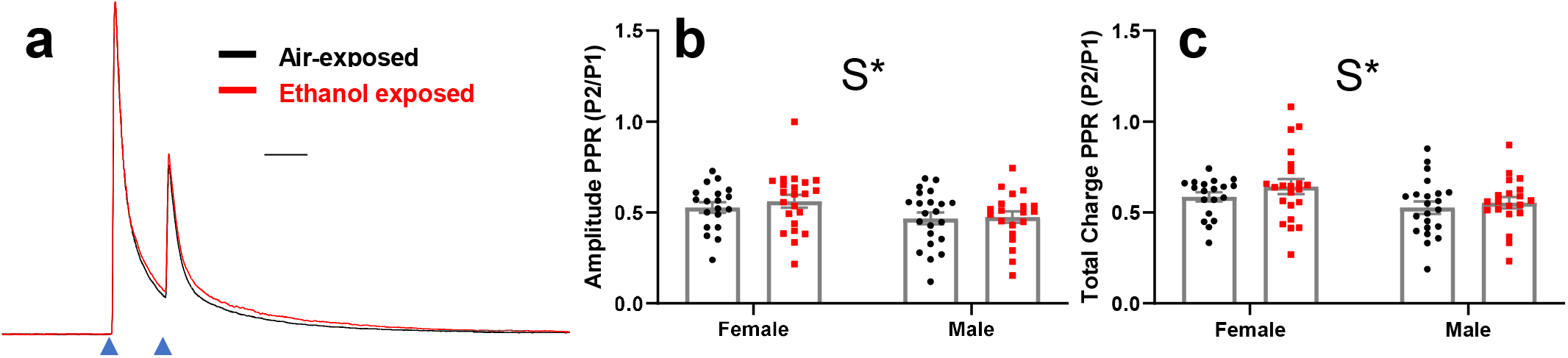
Effect of P7 ethanol exposure on optically-evoked inhibitory postsynaptic currents (oIPSCs) paired-pulse ratios in layer V RSC pyramidal neurons at P40-60. a) Average oIPSC PPR current traces from air-exposed (black trace) and ethanol-exposed (red trace) animals, collapsed across sex. Amplitudes of traces from air- and ethanol-exposed animals are normalized to the amplitude of the first peak to illustrate PPRs. Scale bar = 40 ms. b) Collected amplitude PPRs for all cells from air-exposed (black circles) and ethanol-exposed (red squares) animals. c) Collected total charge PPRs for all cells from air-exposed and ethanol-exposed animals. Asterisk (*) denotes a p-value of p < 0.05. Please see Supplemental Table 2 for exact p-values. S = sex effect. Air female n = 19 cells from 5 animals from 4 litters, air male n = 22 cells from 6 animals from 5 litters; ethanol female n = 22 cells from 6 animals from 5 litters; ethanol male n = 20 cells from 5 animals from 5 litters. Data are presented as mean ± SEM.

## DISCUSSION

The results of our study advance knowledge of the short- and long-term effects of developmental ethanol exposure on GABA_A_ receptor-mediated transmission in the cerebral cortex. We demonstrate that acute ethanol exposure does not affect GABA_A_ receptor function in the RSC of neonatal mice, suggesting that potentiation of these receptors is not involved in the mechanism responsible for the apoptotic neurodegeneration triggered in this brain region by binge-like ethanol exposure during the brain growth spurt. Moreover, we show that this ethanol exposure paradigm causes long-lasting functional alterations at PV-IN→pyramidal neuron synapses from adolescent mice. These findings identify a novel mechanism that could contribute to the cognitive alterations associated with fetal ethanol exposure.

In Experiment #1, we demonstrate that eGABA_A_-PSCs in layer V neurons of the mouse RSC are not affected by acute ethanol exposure during the third trimester-equivalent of human pregnancy. These results follow up on those of a recent study from our laboratory, in which we showed that acute exposure to ethanol during P6-P8 inhibits the excitability of RSC layer V pyramidal neurons and INs mainly through effects exerted post-synaptically on NMDA receptors^28^. Together, the outcomes of these two studies suggest that apoptosis caused by binge-like exposure to ethanol in the RSC during this critical developmental period is caused by inhibition of neuronal activity mediated by a decrease in the function of NMDA receptors, rather than potentiation of Cl^-^ current flow through GABA_A_ receptors. To determine if this lack of an effect of ethanol on eGABA_A_-PSCs was specific to the RSC, we also measured currents in developing CA1 hippocampal pyramidal neurons. We found that acute 90 mM ethanol exposure did increase eGABA_A_-PSC amplitude but for a relatively short duration. A previous study with hippocampal slices from mice showed that protein kinase C (PKC) delta increases, whereas PKC epsilon decreases, sensitivity of IPSCs to acute ethanol (80 mM) exposure (including duration of ethanol’s potentiation)^54^. These PKC isoforms have been demonstrated to be expressed in the hippocampus of P6 mice^55^. Future studies should determine whether the relative activity levels of these kinases in vicinity of GABA_A_ receptors modulate their sensitivity to ethanol during early postnatal development.

One of the hypotheses to explain why ethanol induces apoptotic neurodegeneration during the third trimester equivalent was developed by John Olney and colleagues about 20 years ago, after observing that patterns of cell death in the neonatal brain caused by ethanol exposure were similar to those produced by the noncompetitive NMDA receptor antagonist, MK-801 and the GABA_A_ receptor positive allosteric modulator, phenobarbital^22^. These investigators concluded that ethanol causes neurodegeneration via excessive inhibition of neuronal activity triggered by both NMDA receptor inhibition and GABA_A_ receptor potentiation. Although a number of studies have demonstrated that ethanol can potentiate post-synaptic GABA_A_ receptors under some conditions, there is substantial evidence indicating that the function of these receptors is not directly enhanced by ethanol in many neuronal populations; rather, an increase in GABA release appears to play a central role in the GABA-mimetic effect of ethanol (reviewed in^56^ and^57^). This also appears to be the case for developing neurons during the brain growth spurt, as indicated by electrophysiological studies with slices from neonatal rats showing that acute ethanol increases GABA release while having minimal effects on postsynaptic GABA_A_ receptor function in layer II and III pyramidal neurons of the parietal cortex^58^, pyramidal neurons and INs of the CA3 hippocampal region^59,60^, and hypoglossal motoneurons in the brain stem^61^. In contrast to the findings of these studies, we found that ethanol does not affect the amplitude of electrically evoked GABA_A_-PSCs in RSC pyramidal neurons and INs. As a positive control, we measured the effect of flunitrazepam on eGABA_A_-PSCs and found that this agent prolongs the duration of these events. This finding indicates that the insensitivity of GABA_A_ receptors to ethanol is not due to a general lack of sensitivity of the receptors to allosteric modulators. In addition, ethanol did not modulate the amplitude, decay, or frequency of spontaneous GABA_A_ receptor-mediated PSCs in these cells. These findings suggest that acute ethanol exposure has neither a presynaptic effect nor a postsynaptic effect on GABA_A_ receptor-mediated synaptic transmission at RSC neurons. Future experiments should investigate the reasons for the insensitivity of developing RSC neurons to the facilitatory effects of ethanol on GABA release, which may be due to differences in the properties of presynaptic ion channels, G protein-coupled receptors, and/or intracellular signaling pathways^62,63^. The results of our past and present studies on the effect of acute ethanol exposure support the notion that ethanol-induced cell death in the RSC is most likely triggered by inhibition of postsynaptic NMDA receptor function rather than potentiation of GABA_A_ receptor-mediated neurotransmission either at the presynaptic or postsynaptic levels.

It has been demonstrated that third-trimester equivalent ethanol exposure results in long-term loss of INs in the rodent brain^13,64^, including PV-INs in the RSC^17,65^. In Experiment #2, we demonstrate that binge-like ethanol exposure during this developmental period causes long-lasting alterations in the function of PV-IN→pyramidal neuron synapses. oIPSC amplitude and total charge were increased in ethanol-exposed animals, suggesting that postsynaptic GABA_A_ receptor function is altered in RSC pyramidal neurons. Future studies will be needed to determine the underlying mechanism of this effect (e.g., increases in receptor expression or changes in phosphorylation state). An unexpected finding from this study was that ethanol-exposed animals had an increased number of individual oIPSC events following a laser pulse when compared to air-exposed animals. This observation is similar to electrophysiology experiments in which currents are evoked in the presence of extracellular strontium instead of calcium, which leads to asynchronous presynaptic release of neurotransmitter^66^. In these studies, an increase in the frequency of postsynaptic events following stimulation is indicative an increase in the probability of presynaptic neurotransmitter release^67^. Similar to our study, Pulizzi et al. observed an increasing number of synaptic release events with increasing laser stimulus durations, which they attributed to increases in presynaptic neurotransmitter release probability^52^. Based on these studies, the increased number of oIPSC events in ethanol-exposed mice observed here indicates that P7 ethanol exposure causes a long-term increase in presynaptic function. However, we did not observe a change in PPRs cause by P7 ethanol exposure, suggesting that ethanol exposure did not produce a global change in synchronous and asynchronous GABA release. Asynchronous transmitter release is mediated by different presynaptic proteins (e.g., the slow Ca^2+^ binding sensors synaptotagmin 7 and Doc2) than synchronous release (e.g., the fast Ca^2+^ binding sensor synaptotagmin 1)^53^. It is possible that ethanol exposure selectively increased the expression and/or function of these and other proteins that regulate asynchronous GABA release from PV INs^53^. Another interesting finding from this study concerns the reduction in oIPSC rise time in ethanol exposed animals. Alterations in rise-time can be indicative of changes in post-synaptic GABA_A_ receptor subunit composition^68^, receptor kinetics^69^, GABA diffusion out of synaptic cleft^70^ or synaptic vs. extra-synaptic receptor localization^71^. It is also possible that rise time was affected by dendritic filtering, as rise time increases as a function of distance of the synapse from the somatic recording location^72,73^. Determining the reason for ethanol-induced changes in rise time is an intriguing avenue for further investigation.

Recent studies that have examined the effect of developmental ethanol exposure on the function of INs. Delatour et al.^74^ found that binge-like prenatal ethanol exposure (embryonic days 13.5 and 16.5) increases sIPSC frequency and charge transfer in layer V/VI pyramidal neurons of the somatosensory cortex of adolescent mice. These investigators also found an increase in the optically-evoked charge transfer PPRs for IPSCs mediated by GABA release at PV-IN→pyramidal neuron synapses. The alterations in the charge transfer reported in that study are consistent with those we report here, indicating that this could be a common effect of ethanol exposure during the equivalents to the second and third trimesters of human pregnancy. Moreover, long-lasting disruptions in synaptic transmission at PV-IN→medium spiny neuron synapses in the dorsolateral striatum were detected in mice exposed to ethanol vapor during the equivalent to all trimesters of human pregnancy^40^. Taken together, our results and those of these electrophysiological studies propound the idea that deficits in the function of PV-INs could play a central role in the pathophysiology of FASD.

Deficits in learning and memory processes are among the most common negative consequences of developmental ethanol exposure^19^. Enhancement of postsynaptic GABAergic neurotransmission in the RSC after third-trimester equivalent ethanol exposure could have several impacts on limbic system network activity that may, in part, underlie the learning and memory deficits that have been documented in mice subjected to binge-like ethanol exposure at P7^24,27^. The RSC has reciprocal connections with several brain areas critical to learning and memory functions, including the hippocampal formation, the entorhinal cortex, and anterior/lateral thalamic nuclei^20^. Importantly, communication between the hippocampus and neocortex via sharp wave ripples involves a subiculum-RSC pathway^75^. Consequently, the RSC has a distinct role in spatial memory; it processes distal spatial cues and allows animals to orient themselves both spatially and directionally^76,77^. Lesioning or inactivating the RSC in rodents and macaques impairs performance on spatial memory tasks, while humans with damage to the RSC have problems with anterograde memory formation and navigation using spatial cues^20,78^. The contribution of PV-mediated GABAergic neurotransmission to RSC function and its role in spatial memory are poorly understood at present. However, recent work has demonstrated that PV-IN activity in the anterior cingulate cortex, to which the RSC is immediately posterior, is necessary for memory consolidation^79^. Given that the RSC and the anterior cingulate are adjacent and strongly interconnected^80^, it is reasonable to hypothesize that a disruption in PV-IN-mediated signaling in the RSC will likewise impact learning and memory processes. Future work can test this hypothesis by capitalizing on advances in chemogenetic strategies to determine if up- or down-regulation of PV-IN activity impacts RSC physiology and/or performance on spatial memory tasks^78^.

In conclusion, our findings indicate that third trimester-equivalent ethanol exposure has limited acute effects on postsynaptic GABA_A_ receptor activity in the RSC but has an enduring impact on GABAergic transmission at PV-IN→pyramidal neuron synapses. Long-term alterations in postsynaptic GABA_A_ receptor function caused by apoptosis of PV-INs in the RSC could contribute to some of the behavioral alterations observed in animal models of FASD^27^. Our findings suggest that detailed evaluation of the function of the RSC and related para-hippocampal cortical regions is warranted in humans with FASD.

## ACKNOWLEDGEMENTS

Supported by NIH grants R37 AA015614 and P50 AA022534 (CFV); Undergraduate Pipeline Network Program and Maximizing Access to Research Career Programs (MJB). This research was partially supported by UNM Comprehensive Cancer Center Support Grant NCI P30CA118100 and made use of the Fluorescence Microscopy and Cell Imaging shared resource. We would like to thank Dr. Deidre Hill at the University of New Mexico Clinical & Translational Science Center for an initial consultation regarding the implementation of LMMs (supported by NIH grant UL1TR001449).

## AUTHOR CONTRIBUTIONS

CWB and CFV designed experiments; CWB, GJC, MJB performed experiments; CWB, GJC, MJB analyzed data, CWB and CFV interpreted results and wrote the manuscript.

