## Supplementary Materials for "ENHANCEMENT OF PARVALBUMIN INTERNEURON-MEDIATED NEUROTRANSMISSION IN THE RETROSPLENIAL CORTEX OF ADOLESCENT MICE FOLLOWING THIRD TRIMESTER-EQUIVALENT ETHANOL EXPOSURE"

**This PDF file includes:**
Supplementary Figures 1 to 4
Supplemetary Tables 1 to 3

### Supplemental Figure 1

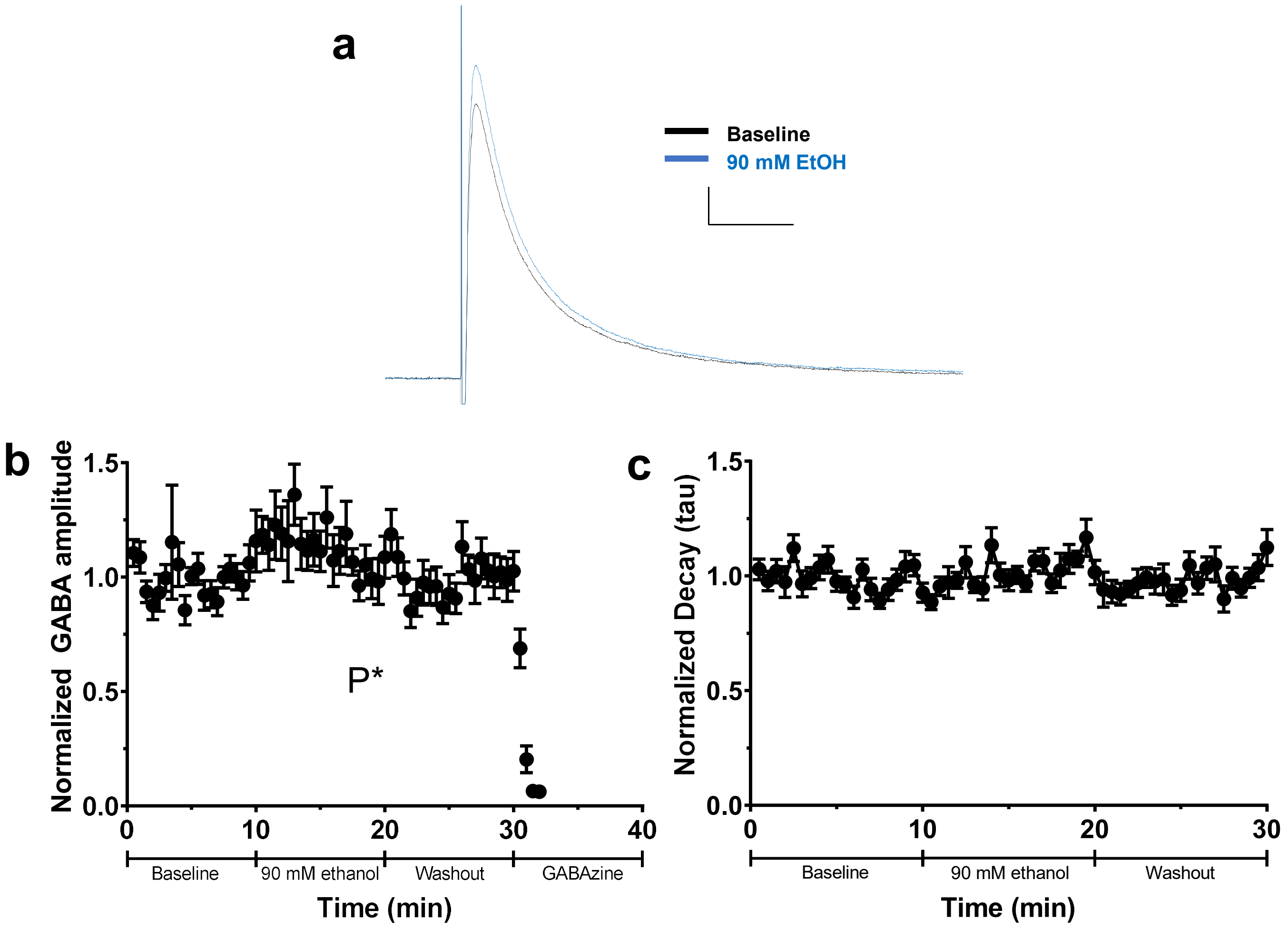


**Supplemental Figure 1. Effect of acute ethanol application on evoked $GABA_A$ receptor-mediated postsynaptic current ($GABA_A$-ePSC) amplitude and decay in CA1 hippocampal pyramidal neurons.** a) Average ePSC traces from CA1 pyramidal neurons during baseline (black trace) and the first 5 minutes of the acute 90 mM ethanol application phase. Scale bars = 40 ms, 20 pA. b) Normalized $GABA_A$-ePSC amplitudes during the baseline, 90 mM ethanol application, washout, and gabazine (25 µM) application phases. c) Normalized decay constants (tau) for CA1 pyramidal neurosn during the baseline, 90 mM ethanol application and washout phases. Data are presented as mean ± SEM. P* denotes a one-way repeated measures ANOVA p-value of 0.0294 for effect of phase comparing the last 5 min of baseline to the first 5 min of the 90 mM ethanol application phase and the first 5 min of the washout phase.

### Supplemental Figure 2

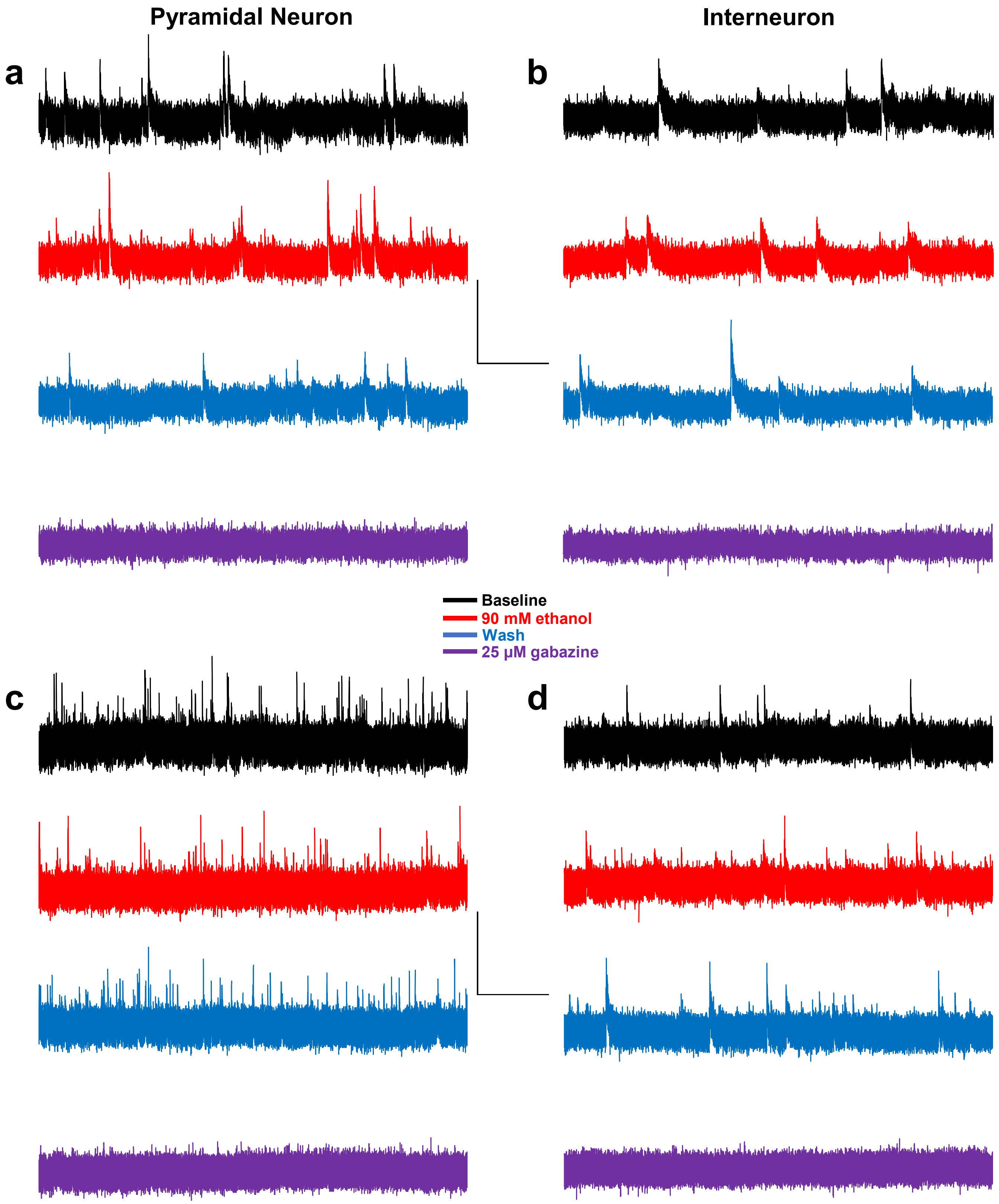

**Supplemental Figure 2. Representative expanded and compressed sPSC traces from a pyramidal neuron and an interneuron demonstrating the effect of 90 mM ethanol application, washout, and 25 µM gabazine application.** a) Expanded traces from a pyramidal neuron during baseline (black trace), 90 mM ethanol application (red trace), washout (blue trace), and 25 µM gabazine application (purple trace). b) expanded traces from an interneuron. Scale bar for expanded traces = 100 ms, 50 pA. c) compressed traces from a pyramidal neuron. d) compressed traces from an interneuron. Scale bar for compressed traces = 5 s, 50 pA.

### Supplemental Figure 3

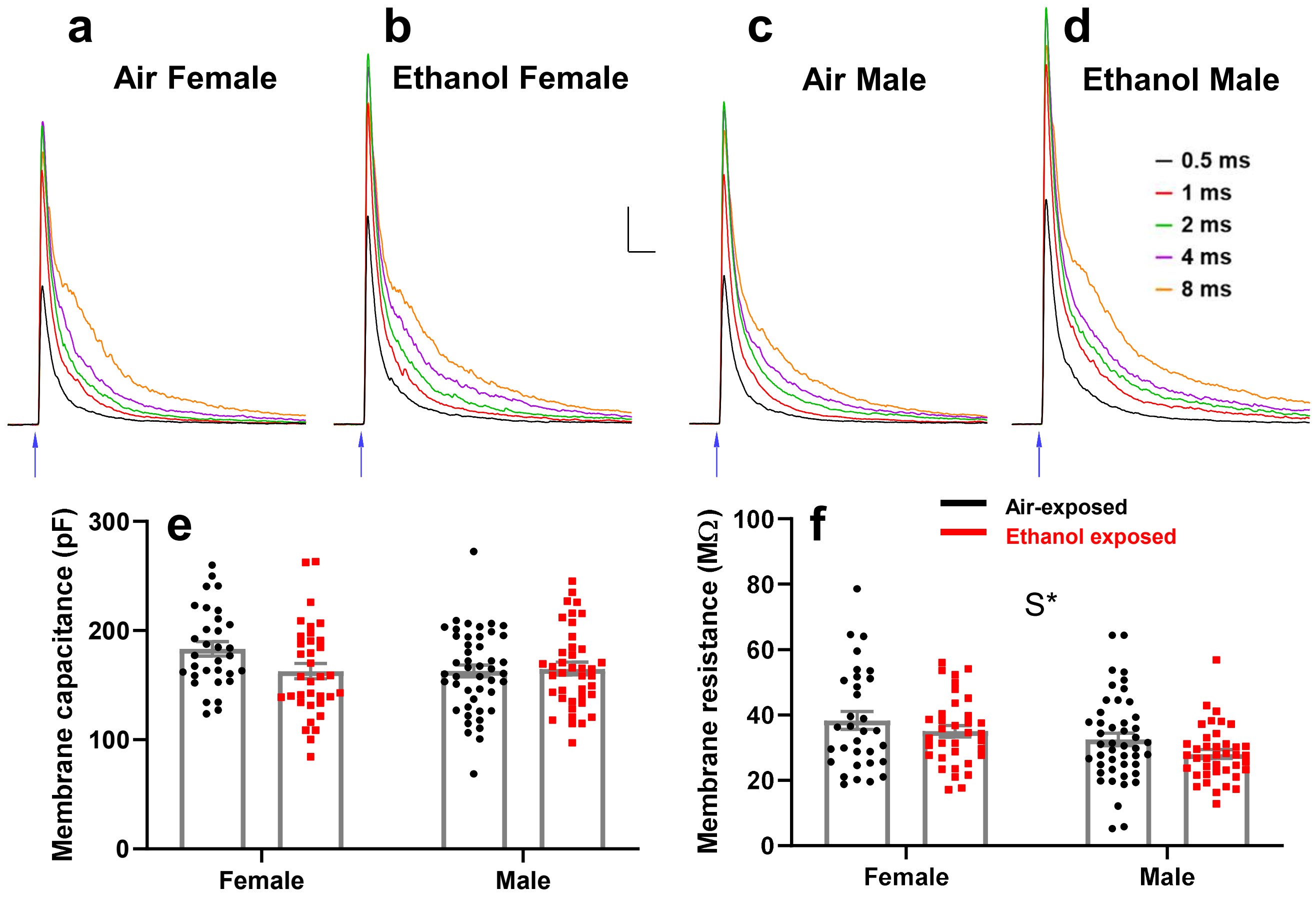


**Supplemental Figure 3. Average optically-evoked inhibitory postsynaptic current (oIPSC) traces for each laser pulse duration in air- and ethanol-exposed animals and analysis of cell membrane capacitance and resistance presented separately by sex.** a-d) Average oIPSC traces from air-exposed female (a), ethanol-exposed female (b), air-exposed male (c), and ethanol-exposed male (d) animals using 0.5 ms (black trace) 1 ms (red trace) 2 ms (green trace) 4 ms (purple trace) and 8 ms (orange trace) laser pulse durations. Scale bars = 20 ms,200 pA. Blue arrows indicate onset of laser pulse e) Membrane capacitances from recordings of optically-evoked inhibitory postsynaptic currents (oIPSC) presented separately by sex. Black circles are values from air-exposed animals, and red squares are values from ethanol-exposed animals. f) Membrane resistances from oIPSC recordings presented separately by sex. Asterisk (*) indicates a p-value of < 0.05. S indicates an effect of sex. Female air n = 32 cells from 8 animals from 7 litters, male air n = 46 cells from 9 animals from 8 litters; female ethanol n = 35 cells from 8 animals from 8 litters, male ethanol n = 40 cells from 8 animals from 7 litters. Data are presented as mean ± SEM.

### Supplemental Figure 4

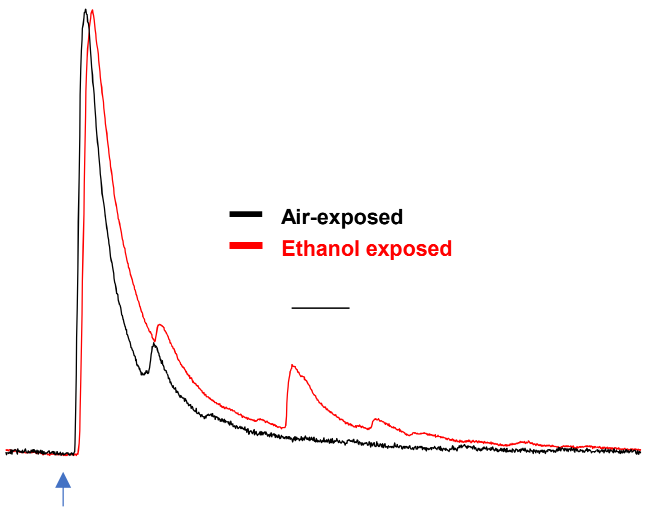


**Supplemental Figure 4. Representative traces demonstrating asynchronous activity evoked by optogenetic stimulation.** Traces show multiple oIPSC peaks evoked from 1 ms laser stimulation (blue arrow shows onset of 1 ms laser stimulation. The black trace is from an air-exposed control animal, the red trace is from an ethanol exposed animal. Traces are normalized by maximal peak amplitude. Scale bar = 10 ms.

| Figure or results section | Experiment | Test | Measure | df | Test statistic | p value | Effect size measure | Effect size | Notes | Passes SW normality test? |
|---|---|---|---|---|---|---|---|---|---|---|
| Figure 2c | Effect of acute ethanol on ePSC amplitude | Repeated measures two-way ANOVA | Interaction: Exposure Phase X Cell Type | F(1.421,22.742) | 0.369 | 0.6237 | Partial eta squared | 0.023 | Did not pass Mauchly's test of sphericity, using Greenhouse-geisser corrected F-ratios and p-values | Yes: residuals pass |
| | | | Main effect: Exposure phase | F(1.421,22.742) | 3.934 | 0.0467 | Partial eta squared | 0.197 | | |
| | | | Main effect: Cell type | F(1,16) | 5.225 | 0.0362 | Partial eta squared | 0.246 | | |
| | | Post-hoc multiple comparison | Baseline vs. 90 mM EtOH | t(17) | 2.470 | 0.0732 | Hedges' g | 0.569 | Bonferroni adjusted p-value | |
| | | | Baseline vs. Wash | t(17) | 2.031 | 0.1746 | Hedges' g | 0.468 | Bonferroni adjusted p-value | |
| | | | 90 mM EtOH vs. Wash | t(17) | 0.322 | >0.9999 | Hedges' g | 0.074 | Bonferroni adjusted p-value | |
| Figure 2d | Effect of acute ethanol on ePSC decay (tau) | Friedman's ANOVA | Main effect: Exposure phase | $\chi^2(2)$ | 2.111 | 0.3480 | Kendall's W | 0.059 | | No: residuals fail normality test |
| | | Mann-Whitney U | Main effect: Cell type | U(n1 = 10, n2 = 8) | 25 | 0.2031 | r | 0.314 | | |
| Figure 3c | Effect of acute flunitrazepam on ePSC amplitude | Repeated measures two-way ANOVA | Interaction: Exposure Phase X Cell Type | F(1.235,32.117) | 0.376 | 0.5885 | Partial eta squared | 0.014 | Did not pass Mauchly's test of sphericity, using Greenhouse-geisser corrected F-ratios and p-values | Yes: residuals pass |
| | | | Main effect: Exposure phase | F(1.235,32.117) | 2.000 | 0.1648 | Partial eta squared | 0.071 | | |
| | | | Main effect: Cell Type | F(1,26) | 11.562 | 0.0022 | Partial eta squared | 0.308 | | |
| Figure 3d | Effect of acute flunitrazepam on ePSC decay | Friedman's ANOVA | Main effect: Exposure phase | $\chi^2(2)$ | 39.071 | <0.0001 | Kendall's W | 0.698 | | No: residuals fail normality test |
| | | Mann-Whitney U | Main effect: Cell type | U(n1 = n2 = 14) | 62 | 0.1035 | r | 0.313 | | |
| | | Friedman's ANOVA post-hoc multiple comparisons (Dunn's multiple comparisons test) | Baseline vs. 1 μM flunitrazepam | U(n1 = n2 = 28) | z = 5.479 | <0.0001 | r | 1.035 | Bonferroni adjusted p-value | |
| | | | Baseline vs. Wash | U(n1 = n2 = 28) | z = 5.345 | <0.0001 | r | 1.010 | Bonferroni adjusted p-value | |
| | | | 1 μM flunitrazepam vs. Wash | U(n1 = n2 = 28) | z = 0.134 | >0.9999 | r | 0.025 | Bonferroni adjusted p-value | |
| Figure 4c&d | Effect of acute ethanol on sPSC frequency | Repeated measures two-way ANOVA | Interaction: Exposure Phase X Cell Type | F(1.382, 17.969) | 0.345 | 0.6344 | Partial eta squared | 0.026 | Did not pass Mauchly's test of sphericity, using Greenhouse-geisser corrected F-ratios and p-values | Yes: residuals pass |
| | | | Main effect: Exposure phase | F(1.382, 17.969) | 3.485 | 0.0670 | Partial eta squared | 0.211 | | |
| | | | Main effect: cell type | F(1,13) | 9.980 | 0.0075 | Partial eta squared | 0.434 | | |
| Figure 4e&f | Effect of acute ethanol on sPSC amplitude | Repeated measures two-way ANOVA | Interaction: Exposure Phase X Cell Type | F(1.336, 17.371) | 6.108 | 0.0173 | Partial eta squared | 0.320 | Did not pass Mauchly's test of sphericity, using Greenhouse-geisser corrected F-ratios and p-values | Yes: residuals pass |
| | | | Main effect: Exposure phase | F(1.336, 17.371) | 4.913 | 0.0314 | Partial eta squared | 0.274 | | |
| | | | Main effect: cell type | F(1,13) | 0.137 | 0.7173 | Partial eta squared | 0.010 | | |
| | | Post-hoc multiple comparison: within pyramidal neurons | Baseline vs. 90 mM EtOH | t(7) | 1.381 | 0.6295 | Hedges' g | 0.274 | Bonferroni adjusted p-value | |
| | | | Baseline vs. Wash | t(7) | 1.373 | 0.6362 | Hedges' g | 0.192 | Bonferroni adjusted p-value | |
| | | | 90 mM EtOH vs. Wash | t(7) | 1.029 | >0.9999 | Hedges' g | 0.081 | Bonferroni adjusted p-value | |
| | | Post-hoc multiple comparison: within interneurons | Baseline vs. 90 mM EtOH | t(6) | 1.110 | 0.9283 | Hedges' g | 0.216 | Bonferroni adjusted p-value | |
| | | | Baseline vs. Wash | t(6) | 2.357 | 0.1695 | Hedges' g | 0.856 | Bonferroni adjusted p-value | |
| | | | 90 mM EtOH vs. Wash | t(6) | 2.234 | 0.2007 | Hedges' g | 0.642 | Bonferroni adjusted p-value | |
| Figure 4g&h | Effect of acute ethanol on sPSC rise-time | Repeated measures two-way ANOVA | Interaction: Exposure Phase X Cell Type | F(2,26) | 0.265 | 0.769524 | Partial eta squared | 0.020 | Passed Mauchly's test of sphericity | Yes: residuals pass |
| | | | Main effect: Exposure phase | F(2,26) | 2.763 | 0.081654 | Partial eta squared | 0.175 | | |
| | | | Main effect: cell type | F(1,13) | 1.448 | 0.250276 | Partial eta squared | 0.100 | | |
| Figure 4i&j | Effect of acute ethanol on sPSC decay (tau) | Repeated measures two-way ANOVA | Interaction: Exposure Phase X Cell Type | F(2,24) | 0.123 | 0.885017 | Partial eta squared | 0.010 | Passed Mauchly's test of sphericity; Removed one interneuron with outlier wash decay value | Yes: residuals pass |
| | | | Main effect: Exposure phase | F(2,24) | 0.834 | 0.446609 | Partial eta squared | 0.065 | | |
| | | | Main effect: cell type | F(1,12) | 13.543 | 0.003148 | Partial eta squared | 0.530 | | |
| Figure 5e | % Colocalization PV-tdTomato/ChR2 | Mann-Whitney U | Effect of P7 ethanol exposure | U(n1 = n2 = 5) | 12 | >0.9999 | r | 0.047 | | No |
| Figure 5f | % Nonspecific transgene expresion | Unpaired t-test | Effect of P7 ethanol exposure | t(8) | 0.406 | 0.6954 | Hedges' g | 0.232 | | Yes |
| Supplemental figure 1b | Effect of acute ethanol on ePSC amplitude (CA1 pyramidal neurons) | One-way ANOVA | Main effect: Exposure phase | F(2,24) | 4.101 | 0.0294 | Partial eta squared | 0.255 | Passed Mauchly's test of sphericity. Baseline is compared to first 5 minutes of EtOH and wash phases. Bonferroni corrected p-values for muliple comparisons | Yes: residuals pass |
| | | Post-hoc multiple comparison | Baseline vs. 90 mM EtOH | t(12) | 2.258 | 0.1299 | Hedges' g | 0.396 | | |
| | | | Baseline vs. Wash | t(12) | 0.663 | >0.9999 | Hedges' g | 0.106 | | |
| | | | 90 mM EtOH vs. Wash | t(12) | 2.405 | 0.0996 | Hedges' g | 0.481 | | |
| Supplemental figure 1c | Effect of acute ethanol on ePSC decay (tau, CA1 pyramidal neurons) | One-way ANOVA | Main effect: Exposure phase | F(2,24) | 0.484 | 0.6224 | Partial eta squared | 0.039 | Passed sphericity test. Baseline compared to first 5 minutes of EtOH and wash phases | Yes: residuals pass |

**Supplemental Table 1: Comprehensive collection of statistical analyses for Experiment 1 and IHC analyses from Experiment 2.**
Detailed information regarding specific tests used, measures examined, degrees of freedom, test statistics, p-values, effect sizes, and results of normality testing are presented.

| Figure | Experiment | Does random effect of litter significantly improve LMM? | Do heterogenous error variances significantly improve LMM? | Exposure | Sex | Laser | Exposure*Sex | Exposure*Laser | Sex*Laser | Exposure*Sex*Laser | Mann-Whitney U: Exposure | Mann-Whitney U: Sex | Notes |
|---|---|---|---|---|---|---|---|---|---|---|---|---|---|
| Supplemental Figure 3e | Effect of P7 ethanol exposure on oIPSC cell capacitance | No | No | F(1,149) = 2.137<br>p = 0.1459<br>*g* = 0.186 | F(1,149) = 2.089<br>p = 0.1505<br>*g* = 0.221 | - | F(1,149) = 3.274<br>p = 0.0724 | - | - | - | - | - | |
| Supplemental figure 3f | Effect of P7 ethanol exposure on oIPSC membrane resistance | Yes | Yes | F(1,19.644) = 0.763<br>p = 0.3930<br>*g* = 0.288 | F(1,95.267) = 4.199<br>p = 0.0432<br>*g* = 0.498 | - | F(1,95.267) = 0.004<br>p = 0.9471 | - | - | - | U(n1 = 78 n2 = 75) = 2550<br>p = 0.1711<br>*r* = 0.111 | U(n1 = 67, n2 = 86) = 2094<br>p = 0.0038<br>*r* = 0.234 | |
| Figure 6c | Effect of P7 ethanol exposure on oIPSC amplitude | No | Yes | F(1,146.828) = 5.009<br>p = 0.0267<br>*g* = 0.328 | F(1,146.828) = 0.518<br>p = 0.4730<br>*g* = 0.075 | F(4,138.866) = 73.641<br>p < 0.0001 | F(1,146.828) = 0.082<br>p = 0.7745 | F(4,138.866) = 1.012<br>p = 0.4035 | F(1,138.866) = 1.015<br>p = 0.4019 | F(4,138.866) = 0.712<br>p = 0.5850 | U(n1 = 387,n2 = 372) = 57494<br>p < 0.0001<br>*r* = 0.174 | U(n1 = 333, n2 = 426) = 67014<br>p = 0.1915<br>*r* = 0.047 | |
| Figure 6d | Effect of P7 ethanol exposure on oIPSC current density | Yes | No | F(1,40.564) = 4.044<br>p = 0.051<br>*g* = 0.384 | F(1,108.138) = 0.687<br>p = 0.4089<br>*g* = 0.157 | F(4,38.243) = 66.409<br>p < 0.0001 | F(1,108.138) = 0.202<br>p = 0.6542 | F(4,38.243) = 1.283<br>p = 0.2936 | F(4,96.088) = 0.816<br>p = 0.5179 | F(4,96.088) = 0.923<br>p = 0.4538 | U(n1 = 386, n2 = 372) = 54471<br>p < 0.0001<br>*r* = 0.209 | U(n1 = 332, n2 = 426) = 64027<br>p = 0.0253<br>*r* = 0.081 | |
| Figure 6e | Effect of P7 ethanol exposure on oIPSC charge | Yes | Yes | F(1,20.848) = 5.907<br>p = 0.0242<br>*g* = 0.424 | F(1,62.016) = 0.848<br>p = 0.3606<br>*g* = 0.123 | F(4,88.351) = 133.734<br>p < 0.0001 | F(1,62.016) = 0.682<br>p = 0.4121 | F(4,88.351) = 2.849<br>p = 0.0284 | F(4,292.749) = 0.682<br>p = 0.6047 | F(4,292.749) = 1.401<br>p = 0.2338 | U(n1 = 381, n2 = 368) = 52361<br>p < 0.0001<br>*r* = 0.219 | U(n1 = 328, n2 = 421) = 63326<br>p = .0516<br>*r* = 0.071 | Compound symmetry covariance matrix for repeated measure residuals |
| Figure 6f | Effect of P7 ethanol exposure on oIPSC half-width | Yes | Yes | F(1,26.478) = 2.315<br>p = 0.1400<br>*g* = 0.270 | F(1,89.647)<br>p = 0.3165<br>*g* = 0.091 | F(4,40.164) = 51.786<br>p < 0.0001 | F(1,89.647) = 0.736<br>p = 0.3932 | F(4,40.164) = 0.604<br>p = 0.6619 | F(4,84.740) = 1.136<br>p = 0.3451 | F(4,84.740) = 0.496<br>p = 0.7389 | U(n1 = 383, n2 = 370) = 60535<br>p = 0.0005<br>*r* = 0.126 | U(n1 = 331, n2 = 422) = 64263<br>p = .0597<br>r = 0.069 | |
| Figure 6g | Effect of P7 ethanol exposure on oIPSC rise time | No | Yes | F(1,110.583) = 11.906<br>p = 0.0008<br>*g* = 0.359 | F(1,110.583) = 0.397<br>p = 0.5297<br>*g* = 0.119 | F(4,102.143) = 15.353<br>p < 0.0001 | F(1,110.583) = 0.713<br>p = 0.4001 | F(4,102.143) = 4.142<br>p = 0.0038 | F(4,102.143) = 1.811<br>p = 0.1324 | F(4,102.143) = 0.567<br>p = 0.6870 | U(n1 = 349, n2 = 324) = 47028<br>p = 0.0002<br>*r* = 0.145 | U(n1 = 288, n2 = 385) = 48535<br>p = 0.0057<br>*r* = 0.107 | |
| Figure 6h | Effect of P7 ethanol exposure on oIPSC asynchronous activity | No | Yes | F(1,137.439) = 13.368<br>p = 0.0004<br>*g* = 0.387 | F(1,137.439) = 0.706<br>p = 0.4020<br>*g* = 0.066 | F(4,133.965) = 126.460<br>p < 0.0001 | F(1,137.439) = 1.804<br>p = 0.1815 | F(4,133.965) = 3.519<br>p = 0.0091 | F(4,133.965) = 0.755<br>p = 0.5564 | F(4,133.965) = 0.563<br>p = 0.6901 | U(n1 = 373, n2 = 371) = 54031.5<br>p < 0.0001<br>*r* = 0.190 | U(n1 = 324, n2 = 420) = 64434.5<br>p = 0.2135<br>*r* = 0.046 | |
| Figure 7b | Effect of P7 ethanol exposure on oIPSC PPR amplitude | No | No | F(1,79) = 0.377<br>p = 0.5408<br>*g* = 0.159 | F(1,79) = 5.119<br>p = 0.0264<br>*g* = 0.507 | - | F(1,79) = 0.169<br>p = 0.6821 | - | - | | - | - | |
| Figure 7c | Effect of P7 ethanol exposure on oIPSC PPR total charge | No | No | F(1,79) = 1.521<br>p = 0.2211<br>*g* = 0.291 | F(1,79) = 5.667<br>p = 0.0338<br>*g* = 0.489 | - | F(1,79) = 0.182<br>p = 0.6709 | - | - | | - | - | |

**Supplemental Table 2: Results from linear mixed-model analyses performed in Experiment 2.**

Details from LMM model building are presented, indicating if including random effect of litter or heterogeneous error variances for vapor chamber exposure conditions significantly improved LMM. F-ratios and p-values are presented for vapor chamber exposure effects, sex effects, repeated measure laser pulse duration effects, and two- and three-way interactions between these effects. Hedges' *g* effect sizes are presented for exposure and sex effects. Non-Parametric Mann-Whitney U tests are presented (including effect size as *r*) for exposure and sex effects from any LMM with residuals that did not pass (p > 0.05) a Shapiro-Wilkes normality test.

| Figure or results section | Experiment | Test statistic and degrees of freedom | p-value | Effect size (Hedges' *g*) | Mann-Whitney U: exposure | Notes |
|---|---|---|---|---|---|---|
| Figure 6e | Effect of P7 ethanol exposure on oIPSC charge: 0.5 msec laser pulse | t(35) = 0.396 | > 0.99 | 0.680 | U(n1 = 73, n2 = 73) = 1793, p = 0.0032, r = 0.282 | Bonferroni corrected p-value |
| | Effect of P7 ethanol exposure on oIPSC charge: 1 msec laser pulse | t(34) = 1.922 | 0.3151 | 0.595 | U(n1 = 77, n2 = 74) = 1947, p = 0.0056, r = 0.265 | Bonferroni corrected p-value |
| | Effect of P7 ethanol exposure on oIPSC charge: 2 msec laser pulse | t(34) = 1.981 | 0.2782 | 0.455 | U(n1 =77, n2 = 74) = 2091, p = 0.0239, r = 0.230 | Bonferroni corrected p-value |
| | Effect of P7 ethanol exposure on oIPSC charge: 4 msec laser pulse | t(34) = 2.526 | 0.0814 | 0.548 | U(n1 = 76, n2 = 73) = 1882, p = 0.0035, r = 0.277 | Bonferroni corrected p-value |
| | Effect of P7 ethanol exposure on oIPSC charge: 8 msec laser pulse | t(34) = 3.363 | 0.0096 | 0.493 | U(n1 = 78, n2 = 74) = 2069, p = 0.0130, r = 0.244 | Bonferroni corrected p-value |
| Figure 6g | Effect of P7 ethanol exposure on oIPSC rise time: 0.5 msec laser pulse | t(107) = 3.472 | 0.0037 | 0.570 | U(n1 = 75, n2 = 66) = 1926, p = 0.1165, r = 0.191 | Bonferroni corrected p-value |
| | Effect of P7 ethanol exposure on oIPSC rise time: 1 msec laser pulse | t(105) = 3.783 | 0.0013 | 0.540 | U(n1 = 72, n2 = 68) = 1881, p = 0.0904, r = 0.200 | Bonferroni corrected p-value |
| | Effect of P7 ethanol exposure on oIPSC rise time: 2 msec laser pulse | t(124) = 0.946 | > 0.99 | 0.215 | U(n1 = 70, n2 = 65) = 1988, p > 0.99, r = 0.109 | Bonferroni corrected p-value |
| | Effect of P7 ethanol exposure on oIPSC rise time: 4 msec laser pulse | t(110) = 2.635 | 0.0482 | 0.390 | U(n1 = 67, n2 = 63) = 1730, p = 0.3815, r = 0.156 | Bonferroni corrected p-value |
| | Effect of P7 ethanol exposure on oIPSC rise time: 8 msec laser pulse | t(110) = 2.635 | 0.0656 | 0.351 | U(n1 = 65, n2= 62) = 1752, p > 0.99, r = 0.113 | Bonferroni corrected p-value |
| Figure 6h | Effect of P7 ethanol exposure on oIPSC asynchronous activity: 0.5 msec laser pulse | t(109) = 3.381 | 0.0050 | 0.565 | U(n1 = 72, n2 = 71) = 1819.5 p = 0.0120, r = 0.254 | Bonferroni corrected p-value |
| | Effect of P7 ethanol exposure on oIPSC asynchronous activity: 1 msec laser pulse | t(113) = 4.322 | 0.0002 | 0.819 | U(n1 = 71, n2 = 75) = 1659.5 p = 0.0004, r = 0.328 | Bonferroni corrected p-value |
| | Effect of P7 ethanol exposure on oIPSC asynchronous activity: 2 msec laser pulse | t(123) = 3.620 | 0.0021 | 0.663 | U(n1 = 74, n2 = 75) = 1921 p = 0.0057, r = 0.266 | Bonferroni corrected p-value |
| | Effect of P7 ethanol exposure on oIPSC asynchronous activity: 4 msec laser pulse | t(144) = 3.191 | 0.0087 | 0.532 | U(n1 = 78, n2 = 75) = 2096 p = 0.0122, r = 0.245 | Bonferroni corrected p-value |
| | Effect of P7 ethanol exposure on oIPSC asynchronous activity: 8 msec laser pulse | t(149) = 1.922 | 0.2824 | 0.337 | (U(n1 = 78, n2 = 75) = 2311 p = 0.1247, r = 0.181 | Bonferroni corrected p-value |

**Supplemental Table 3: Post hoc examination of exposure by laser interactions with a p-value of < 0.05 from LMM analyses.**
Non-parametric Mann-Whitney U tests for exposure effects within each laser pulse duration are presented for the effect of ethanol exposure on oIPSC total charge, oIPSC rise time, and oIPSC asynchronous activity. Data presented include: results of parametric post-hoc tests (not discussed in text as data violated normality assumptions), Mann-Whitney U test statistics with group sample sizes, p-values, and effect sizes as *r*. All p-values are Bonferroni corrected for the number of multiple comparisons made.
